# Machine learning uncovers circulating biomarkers and molecular heterogeneity in obesity and type 2 diabetes

**DOI:** 10.64898/2026.04.16.718836

**Authors:** Erdenetsetseg Nokhoijav, Miklós Káplár, Sándor Csaba Aranyi, András Berzi, Göran Bergström, Konstantinos Antonopoulos, Fredrik Edfors, Miklós Emri, Éva Csősz

## Abstract

Obesity and Type 2 Diabetes (T2D) are heterogeneous metabolic disorders whose molecular diversity is incompletely defined. We analyzed circulating proteomic profiles from 129 individuals belonging to Control, Obesity, and T2D groups and applied complementary machine-learning approaches, including random forest, multinomial logistic regression with LASSO regularization, support vector machines, and ensemble voting to identify proteins distinguishing the clinical groups. Convergent model outputs revealed a partially overlapping panel of discriminative proteins. Model performance was evaluated in an independent dataset from the Human Protein Atlas (n=834) comprising healthy individuals, patients with Obesity, T2D, or other metabolic diseases. Unsupervised clustering further identified multiple proteomic subgroups within each clinical category, indicating substantial intragroup heterogeneity. Bootstrap random forest with null-model benchmarking highlighted stable cluster-discriminative proteins. These findings demonstrate that integrating circulating proteomics with machine learning can resolve molecular heterogeneity in Obesity and T2D and nominate candidate biomarkers for metabolic disease stratification.

## 1. Introduction

Obesity and Type 2 Diabetes (T2D) are among the most prevalent chronic conditions worldwide, imposing substantial public health and economic burden^1^. Both disorders are tightly linked to metabolic dysregulation^2–4^, systemic inflammation^5–7^, and increased risk of cardiovascular and other comorbidities^8,9^. Although Obesity and T2D share overlapping risk factors and pathophysiological pathways, they display considerable heterogeneity in disease onset, progression, and clinical outcomes. This diversity complicates diagnosis, risk stratification, and therapeutic interventions. A deeper molecular understanding of these conditions is therefore essential for identifying biomarkers that can capture subgroup-specific signatures and guide personalized approaches to management.

Proteomics offers powerful and comprehensive approaches for characterizing systemic alterations in such complex diseases. Blood circulating proteins reflect both genetic and environmental influences and may provide direct insights into metabolic pathways altered in Obesity and T2D. Recent advances in high-throughput proteomics platforms, including MS-based methods and affinity-based^10^ assays have enabled quantification of hundreds to thousands of proteins from limited biological materials. However, challenges remain in integrating data across platforms, accounting for redundancies, and identifying the most informative features in high-dimensional datasets. Furthermore, while proteomics studies have revealed numerous candidate biomarkers associated with Obesity and T2D, less is known about the molecular signatures that define heterogeneity to discriminate between these two metabolic conditions as well as within each of these groups.

To address these challenges, we applied different platforms, including Tandem Mass Tag (TMT) labeling with Data-Dependent Acquisition (DDA) method, label-free Data-Independent Acquisition (DIA) analysis^11^, and Olink-Proximity Extension Assay (PEA)^10^ technology. These platforms enabled comprehensive profiling of serum proteins across Controls without Obesity and with normal blood glucose level, individuals with Obesity (BMI>30, no signs of diabetes), and individuals with T2D. By prioritizing non-redundant protein features across platforms, we assembled a dataset suitable for downstream statistical and machine learning (ML) analyses. Our objectives were to identify discriminative proteins across groups using different supervised ML strategies and in parallel, to explore intragroup heterogeneity by applying unsupervised cluster analysis, followed by supervised ML analysis.

## 2. Materials and methods

### 2.1. Study Population

Initially, 187 subjects were recruited for this study^11–13^. Based on the data completeness, 129 participants were included in the ML analysis: 51 patients with T2D, 48 subjects with Obesity, and 30 healthy volunteers.

Patients with T2D were diagnosed by an oral glucose tolerance test (OGTT), regardless of body mass index (BMI). Subjects with Obesity were included if their BMI was ≥ 30 kg/m² and they had no T2D diagnosis at the time of their visit to the Internal Medicine Clinic, University of Debrecen. Healthy Controls were selected based on normal fasting blood glucose levels and BMI and were matched to the T2D and Obesity groups for age and sex.

The median age of participants was 52 (IQR: 45.00–57.00) years for the T2D group, 52 (IQR: 46.75–61.00) years for the Obesity group, and 51 (IQR: 41.50–57.75) years for the Control group. Baseline characteristics are summarized in Table 1.

**Table 1.** Baseline characteristics of the study population.

| Parameters, units | Control<br>(N=30) | Obesity<br>(N=48) | T2D<br>(N=51) | Adj.p-value |
| --- | --- | --- | --- | --- |
| **Age**, years<br>(median [IQR]) | 51.00<br>[41.50–57.75] | 52.00<br>[46.75–61.00] | 52.00<br>[45.00–57.00] | 0.509<sup>a</sup> |
| **Glucose**, mmol/L<br>(median [IQR]) | 4.90<br>[4.70–5.30] | 5.40<br>[5.00–5.80] | 8.30<br>[6.40–9.30] | <0.0001<sup>a</sup> |
| **BMI**, kg/m<sup>2</sup><br>(median [IQR]) | 26.12<br>[25.13–27.10] | 36.32<br>[33.30–40.37] | 32.41<br>[29.36–36.51] | <0.0001<sup>a</sup> |
| **Sex**, N (%) | F: 13 (43.3%);<br>M: 17 (56.7%) | F: 28 (58.3%);<br>M: 20 (41.7%) | F: 20 (39.2%);<br>M: 31 (60.8%) | 0.216<sup>b</sup> |

BMI, Body Mass Index; IQR, Interquartile Range; M, Male; F, Female. Adjusted p values indicate statistical significance ( $p < 0.05$ ) based on <sup>a</sup>Kruskal-Wallis rank-sum test or <sup>b</sup>Pearson's Chi-squared test with Benjamini-Hochberg adjustment.

All participants provided written informed consent according to the Declaration of Helsinki and the study was approved by the Ethics Committee of the University of Debrecen (4845B-2017) and the National Institute of Pharmacy and Nutrition (OGYEI/2829/2017, Approval date: 31 January 2017).

### 2.2. Sample collection

Peripheral blood samples from overnight fasted participants were collected in native tubes, and the serum samples were aliquoted and stored at −80°C until processed by TMT-based, DIA-based analyses^11^, and PEA^12^ analysis.

### 2.3. Clinical assessment

Clinical laboratory and physical examinations were carried out at the Laboratory Medicine Institute and Department of Internal Medicine, University of Debrecen. In total, 61 clinical parameters were used in our study, including 26 parameters for the clinical laboratory, 25 hematology and coagulation parameters, 6 anthropometric measurements, and several calculated indices. The LiverRisk Score was calculated with an online tool (https://www.liverriskscore.com) using the gender, age, fasting blood glucose level, as well as the activity of glutamate-oxaloacetate transaminase (GOT)/aspartate aminotransferase (AST), glutamate-pyruvate transaminase (GPT)/alanine aminotransferase (ALT), and gamma-glutamyl transferase (GGT), and platelet counts, according to the tool guidelines^14^.

### 2.4. Proteomics data assessment

#### 2.4.1. Data source

Quantitative proteomics profiling was performed on 235 identified proteins of 90 participants from the TMT dataset, whereas in the DIA dataset, we identified 232 proteins in 134 participants^11^. After clearing the overlaps, 200 proteins from the TMT and 52 proteins from the DIA dataset were subjected to further analyses.

In addition, protein abundance in 130 serum samples was measured using the Olink Explore Cardiometabolic panel, performed by Olink Proteomics (Uppsala, Sweden). From this analysis, Normalized Protein Expression (NPX) values for 366 proteins (Olink dataset) were generated based on the intra- and inter-run normalization and log2-scaling^15^. NPX values were centered by subtracting the mean of each variable and scaled by dividing by its standard deviation (SD).

All three proteomics datasets were subsequently integrated with clinical data to generate the full proteomics dataset.

#### 2.4.2. Data preprocessing

The datasets generated by different proteomics methods were merged. In order to eliminate redundancy and overlaps, we applied prioritization. In case of duplication, the data with higher priority was kept, and the others were removed. The TMT dataset was arbitrarily prioritized over the DIA dataset, and the Olink dataset was prioritized over the TMT and DIA datasets, with overlaps defined by exact protein name matches. This approach ensured a non-redundant protein set for downstream analyses. A data completeness analysis was performed on the non-redundant proteins, and we kept those proteins where the missing data was less than 30%. Protein-wise missingness was assessed within each clinical group. All proteins were complete in the Control group, while a partial missingness was observed for two proteins (ACOX1 and SNAP23) in three patients with Obesity, and for one protein (SNAP23) in one participant with T2D. To retain these proteins and maintain a complete data matrix for multivariate analyses, the missing values were replaced with “0.” No additional imputation was applied.

As part of data preprocessing, we introduced a synthetic control feature entitled “Random” generated by random sampling from the empirical value distribution. This feature served as a negative control to establish a baseline for feature importance and to aid in distinguishing biologically meaningful signals from random noise.

### 2.5. Proteomics data analysis

To improve robustness and interpretability, the ML workflow was organized into three analytical stages:

1. identification of proteins discriminating the clinical groups using multiple supervised learning approaches,
2. biological interpretation and external validation of identified candidate biomarkers, and
3. detection of molecular heterogeneity within groups using unsupervised clustering followed by supervised discrimination of clusters.

Within this framework, Random Forest (RF) was considered the primary discovery model, while Least Absolute Shrinkage and Selection Operator (LASSO), Support Vector Machine (SVM), and Voting Ensemble (VE) approaches were applied as complementary methods to confirm feature robustness and reduce model-specific bias.

#### 2.5.1. Application of ML models to identify proteins discriminating between groups

First, we trained a multiclass RF classifier^16,17^ to discriminate the three groups based on all available protein features as predictors. Modeling was performed using the randomForest package^17^. Feature importance was quantified with the Mean Decrease Gini (MDGini) score. The resulting importance was sorted in descending order, and the top 10 proteins with the highest MDGini values were retained for reporting.

To identify discriminative proteins across study groups by the second model, we applied multinomial logistic regression with LASSO regularization^18^ using the glmnet^19^ package. The model was trained on the full dataset with all three groups, Control, Obesity, and T2D as the outcome variables. Regularization tuning was performed using 10-fold cross-validation (CV). The optimal regularization parameter (λ) was selected using the lambda.1se criterion^20^, which is the largest penalty value within one standard error of the minimum cross-validated error. Model coefficients at lambda.1se were extracted for each group, and the proteins with non-zero coefficients were retained as predictive or important features.

To test our third model for evaluating the discriminative capacity of proteomics features, we trained a SVM classifier^21,22^ using the e1071 package^23^. The predictor matrix consisted of all scaled protein features. To identify the most predictive proteins, we applied Recursive Feature Elimination (RFE) with linear SVM^22^. Model performance across subset sizes was visualized to determine the optimal number of predictors.

Validation for all three ML models was carried out by a 5-fold CV method and reported in terms of accuracy and confusion matrices.

To increase robustness and reduce model-specific bias, we applied a VE approach combining the predictions of RF, LASSO, and SVM models. Feature importance within the ensemble was evaluated using “SHapley Additive exPlanations” (SHAP), which quantify the marginal contribution of each feature to model predictions based on cooperative game theory^24,25^.

#### 2.5.2. Verification of potential biomarkers

The potential biomarker panel was verified on Human Protein Atlas (HPA) samples measured on the Olink Explore 1536 platform^26^. Two general population cohorts were included: Impaired Glucose Tolerance and Microbiota (IGTM)^27^ and Swedish CArdioPulmonary BioImage Study (SCAPIS)^28^ and from these, four groups were defined: Obesity, T2D, Healthy controls, and a broader disease Control group named “Other” comprising samples from patients with metabolic dysfunction-associated steatotic liver disease (MASLD), coronary artery calcification, and metabolic syndrome. In the verification dataset 834 donors were in the Healthy Control group (n=549 from IGTM, n=285 from SCAPIS), 75 in T2D group (n=44 from IGTM and n=31 from SCAPIS), 243 in Obesity group (n=184 from IGTM, n=59 from SCAPIS) and 678 in the Other group (n=455 from IGTM, n=223 from SCAPIS). 100 LASSO models were applied using the discriminatory panel for two classification settings: HOT: Healthy vs Obesity vs T2D, and OTC: Obesity vs T2D vs Healthy and Other. Models were trained on IGTM with class balancing and blindly tested on SCAPIS.

#### 2.5.3. Application of ML models to detect molecular heterogeneity within group

Unsupervised k-means clustering within each clinical group was performed. In all three clinical groups, *k = 2* showed the highest silhouette scores and was therefore selected. The unsupervised ML algorithm, k-means clustering^29–31^, was then applied using kmeans() function^32^ in R.

To identify the features discriminating between the k-means-derived, intragroup clusters, we applied RF model^16^, a supervised ML algorithm suitable for classification tasks, with a cluster column as a target for model training. Due to the relatively low number of subjects in the groups, we applied the bootstrapping technique^33^, allowing to repeatedly draw samples with replacement from the source to assess a particular parameter. One thousand bootstrap samplings were performed, and on every sample a RF model was trained with cluster assignment (*k* = 2) as the outcome and protein abundances as predictors. To mitigate known biases of MDGini score^34^ toward continuous and high-variance features, all predictors were standardized prior to modeling, and importance estimates were benchmarked against null distributions derived from shuffled data using the randomForest^16^ package with bootstrap.

To establish a statistical significance threshold, we generated a so-called shuffled dataset where artificially disturbed the connection between features and target column. Briefly, we looped over all features, and at every step we randomly shuffled rows of the table while keeping a target column intact. This shuffled dataset was subjected to 1000 bootstrap RF iterations as above-mentioned to get information about the importance of features that do not have any relevance in predicting a cluster, thus providing a baseline for feature separation. Then we compared the density distributions of MDGini scores from the real and shuffled data analyses. The intersection point between the two distributions was used as a data-driven threshold. Proteins with average importance values greater than this threshold were retained as informative features, while those below were considered random noise.

Additionally, the scale of feature importance can vary from one bootstrap sample to another. In fact, a feature can show high importance; however, be unstable and show up in a few bootstrap samples. To prioritize robust and reproducible discriminators, we further applied a stability-based criterion, retaining proteins that appeared among the top 30 ranked features in at least 70% of bootstrap iterations. This approach emphasizes reproducibility over single-run importance, and those retained proteins were considered as highly stable important features.

The bootstrapped RF model performance was evaluated by mean out-of-bag (OOB) error rates, aggregated confusion matrix, and mean class error per cluster.

### 2.6. Visualization

Intra-group cluster plots were visualized using the factoextra^35^ package in R. The scatter plots display individuals in clusters with each point representing one participant and colored according to its k-means cluster assignment.

The RF results were visualized as scaled dot plots, boxplots, and heatmaps generated using the ggplot2^36^ and ComplexHeatmap^37^ packages in R. The top 30 most important features and the stable features were displayed in scaled dot plots, where the scale size corresponded to either the average MDGini score or the selection frequency across bootstrap iterations. To visualize the discriminative distribution of highly stable important features between clusters, the amounts of those proteins were plotted as boxplots with individual data points. Additionally, the distribution patterns of highly stable important features across clusters and participants were visualized in heatmaps.

### 2.7. Functional analysis

Gene Ontology (GO) enrichment analysis was performed for each set of highly stable important features within the group using the enrichGO function from the R package clusterProfiler^38,39^ with the org.Hs.eg.db^40^ human annotation database (version 3.20.0) in R. Enrichment was the Biological Process (BP), and the p-values were adjusted with Benjamini–Hochberg (BH) method^41^.

### 2.8. Statistical analysis

All statistical and data analysis were performed utilizing R (v4.4.3) statistical programming language^42^ with in-house developed scripts.

Classical non-parametric statistical tests were performed besides the ML-based feature selection. While ML models prioritize features based on predictive contribution, Wilcoxon^43^ and Kruskal-Wallis^44^ tests were applied to assess univariate distributional differences with BH false discovery rate (FDR)^41^ with adjusted p-value < 0.05.

Intergroup and intragroup differences in protein abundance, clinical variables were tested using either Wilcoxon rank-sum test^43^ or Kruskal-Wallis^44^ rank-sum test for each protein with FDR adjustment and significance cut-off was p < 0.05. For the categorical variable, gender, comparisons were made using Pearson’s Chi-square test^45^. All analyses were performed using the tidyverse^46^ and stats^42^ packages. The results were represented as median (IQR) or number (%) as appropriate, and the statistical significances were reported as adjusted p-values in the Supplementary Table 1 and Supplementary Table 2.

## 3. Results

The TMT, DIA, and Olink datasets were subsequently integrated (Supplementary Figure 1), and the extent of missing data was evaluated. Data completeness was high for the Olink dataset, whereas the TMT dataset exhibited a substantial proportion of missing values and was therefore excluded from further analysis. The merged and cleaned dataset consisted of NPX data from 30 Controls, 48 individuals with Obesity, and 51 individuals with T2D (n = 129). One participant with Obesity was excluded due to missing clinical variables other than BMI.

The data analysis followed a three-level strategy: (1) first the proteins making possible the discrimination between the clinical groups were identified followed by their (2) validation on independent datasets, and finally (3) the molecular heterogeneity at the group level was examined and proteins able to define subgroups in the clinically homogenous patient groups were identified.

### 3.1. ML models identify a circulating protein signature distinguishing Obesity and T2D

We applied an integrated approach for data analysis, starting with classical statistical analysis followed by the application of ML to uncover the proteins that can provide a good classification of the study groups.

#### 3.1.1. Classical statistical analysis does not allow for proper discrimination of the study groups

Significantly different proteins between the control, Obesity, and T2D groups were identified using a non-parametric statistical approach. For each protein, the expression values were compared across groups using the Kruskal-Wallis test, followed by FDR correction of p-values (Figure 1).

**Figure 1.**
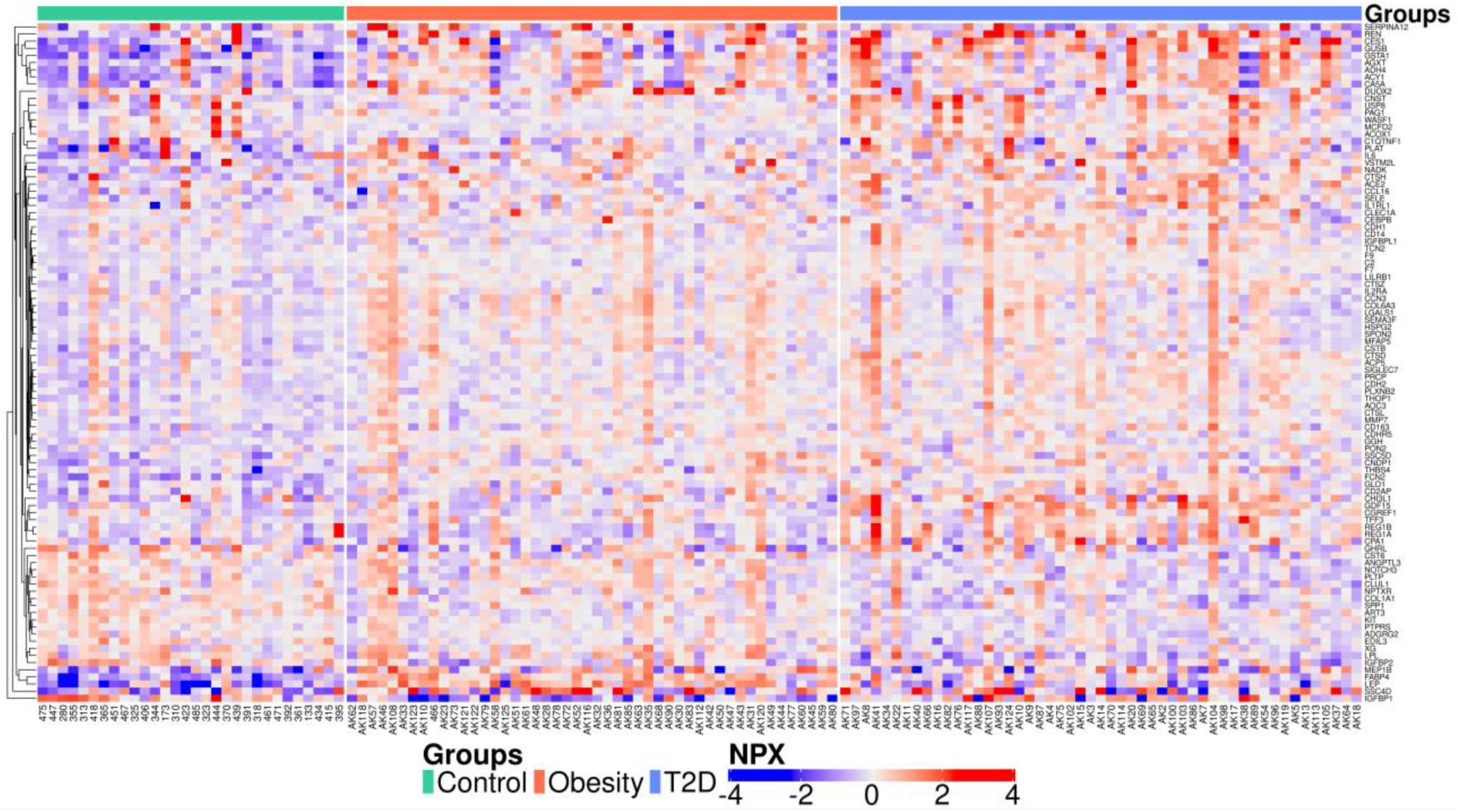
Heatmap of significantly different proteins across groups (N = 95). Rows represent proteins, and columns represent individual participants. Color scale indicates protein NPX values (red = higher, blue = lower). Top column bars denote study groups (green = Control, orange = Obesity, blue = T2D).

95 proteins from our dataset were statistically significantly different between the study groups. The heatmap visualization showed the distinct expression patterns across groups, with good discrimination between control and disease groups, but low discrimination between the Obesity and T2D groups.

#### 3.1.2. ML models revealed group-discriminating proteins

In order to identify proteins able to discriminate the groups from each other, we applied multiclass classifiers including Random Forest (RF), multinomial logistic regression with Least Absolute Shrinkage and Selection Operator (LASSO) regularization^18^, Support Vector Machine (SVM)^21,22^ with Recursive Feature Elimination (RFE), and Voting Ensemble (VE) with feature importance selection by SHapley. The series of results across multiple algorithms increased the robustness of our findings while highlighting the model-specific signature features (Figure 2).

**Figure 2.**
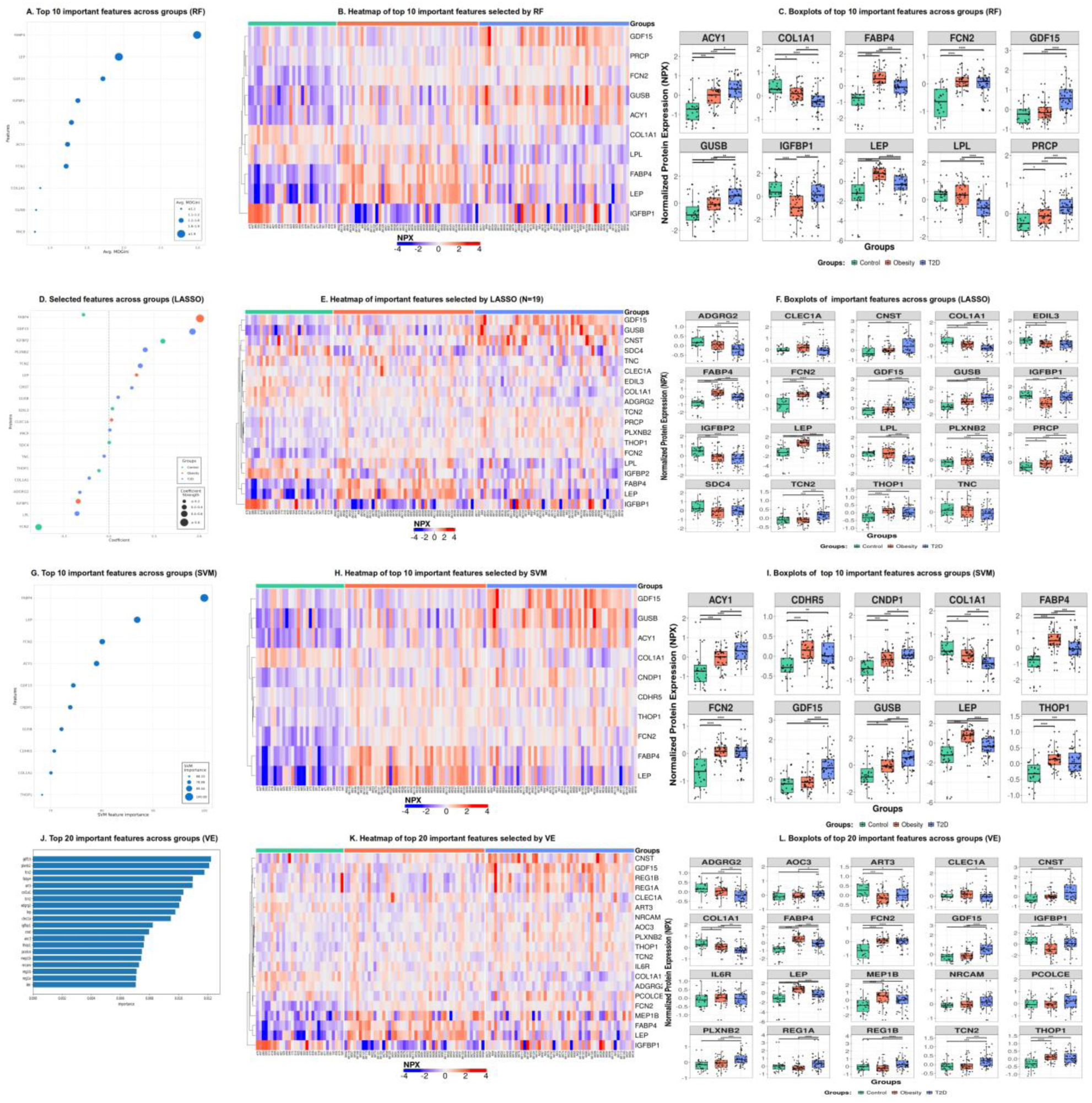
Identification of intergroup discriminating proteins across Control, Obesity, and T2D groups using multiple machine learning classifiers. (A-C) Top 10 important features identified by Random Forest (RF). (D-F) Important features selected by multinomial logistic regression with LASSO (N = 19). (G-I) Top 10 important features identified by SVM. (J-L) Top 20 important features identified by EV. (A, D, G, J) List of selected features by multiple ML models. (B, E, H, K) Heatmaps of selected features by each method, showing protein abundance (NPX) across all participants. Color scale bar represents protein expression levels (purple = low, red = high). (C, F, I, L) Distribution of intergroup discriminating proteins across Control, Obesity, and T2D groups. y axis shows the protein amount across the groups. Asterisks indicate statistically significant differences as adjusted p-values (* adj.p < 0.05, ** adj.p < 0.01, *** adj.p < 0.001).

All above-mentioned ML models^47–50^ are widely used in proteomics studies for multiple diseases.

From the RF model, we selected the top 10 features with the highest MDGini score, including FABP4, LEP, GDF15, IGFBP1, LPL, ACY1, FCN2, COL1A1, GUSB, and PRCP across three groups (Figure 2, panels A, B, C). The LASSO selected a broader set of proteins, 19 features, due to its own selection criterion (Figure 2, panels D, E, F). By SVM model, we identified the top 10 features with CNDP1 and CDHR5 as unique discriminators (Figure 2, panels G, H, I). With VE model, we selected the top 20 features based on their SHAP importance scores. SHAP values allow consistent comparison of feature effects across models and improve the interpretability of complex ML pipelines. Among them, eight features, ART3, AOC3, PCOLCE, MEP1B, NRCAM, REG1B, REG1A, IL6R, were unique discriminators between study groups (Figure 2, panels J, K, L).

Each model prioritized a partly overlapping but distinct set of proteins based on the model-specific selection criteria and listed the important proteins which discriminate the clinical groups from each other.

Heatmaps confirmed that the above-mentioned selected proteins showed consistent expression across groups. Selected proteomic signatures were showing distinctive characteristics for Obesity and T2D groups compared with Control group. Of note, FABP4, LEP, GDF15, FCN2, and COL1A1 were consistently selected by all four models, underscoring their strong discriminatory potential. The boxplots further illustrated the distribution of these selected proteins across Control, Obesity, and T2D groups (Figure 2, panels C, F, I, L). Together, these results demonstrate that while a few core proteins (FABP4, LEP, IGFBP1, COL6A3) robustly discriminate across all methods and groups, additional proteins highlighted by LASSO and SVM may capture complementary biological variation, providing a more nuanced inter-group separation.

#### 3.1.3. Candidate circulating protein biomarkers distinguishing Obesity and T2D

In case of RF model, none of the top 10 features were model-specific; four features, IGFBP2, EDIL3, SDC4, and TNC were uniquely selected by LASSO model; in SVM model, two features, CDHR5 and CNDP1, were unique, while eight, AOC3, ART3, IL6R, MEP1B, NRCAM, PCOLCE, REG1A, REG1B were uniquely selected by VE model, respectively (Figure 3A).

**Figure 3.**
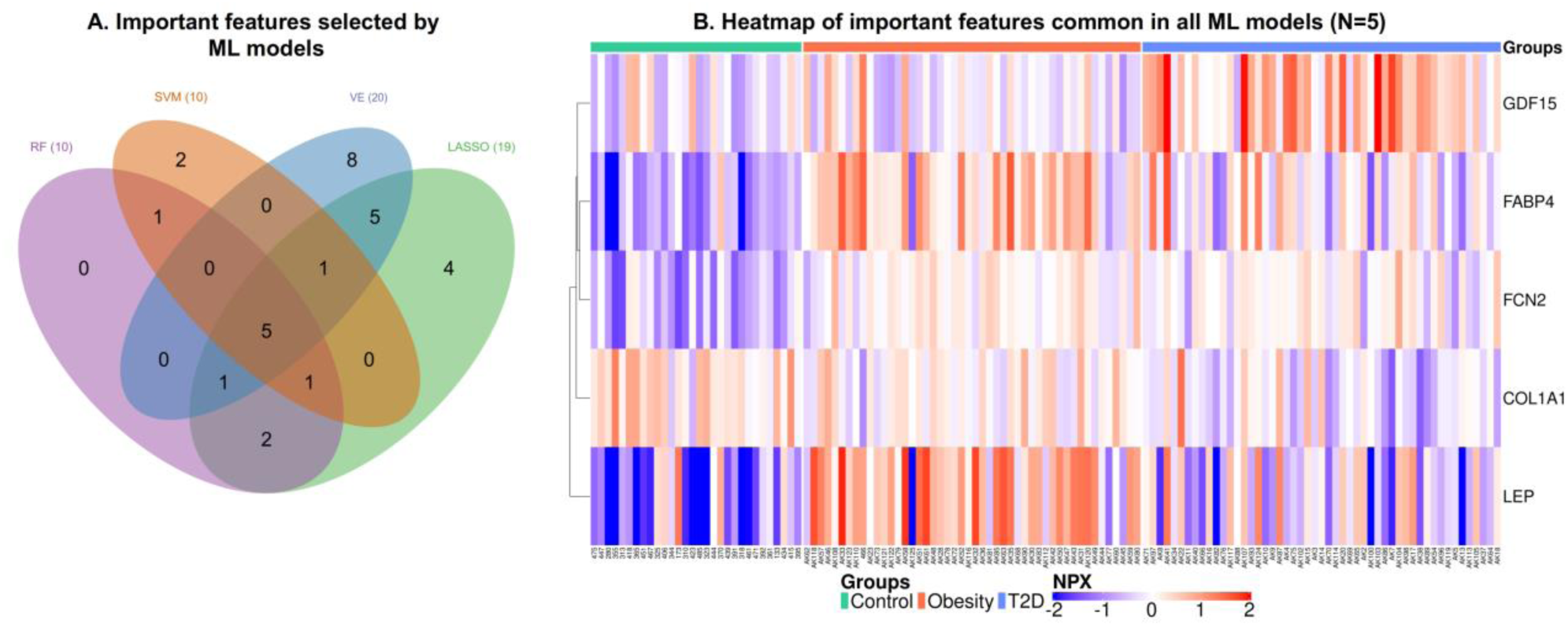
Evaluation of ML models. (A) Venn diagram of features selected by ML models of RF (N = 10), LASSO (N = 19), SVM (N = 10) and VE (N = 20) showing their unique and shared features. (B) Heatmap of five features in common, consistently selected by all four models.

Five features, FABP4, LEP, GDF15, FCN2, and COL1A1, were commonly selected by all models, indicating that they are robust and model-independent group discriminatory features (Figure 3). The performance test on these features was evaluated using a svmRadial method. The dataset was randomly split into a training set (80%) and an independent test set (20%) with stratification by groups to preserve class proportions. Prediction performance was quantified using overall accuracy, where the sum of correctly classified samples was divided by the total number of observations, and the accuracy was: 0.8. Therefore, we used this score as a threshold to select candidate biomarkers. In addition to these five features, we included GUSB to our candidate biomarkers list based on its distinct expression pattern between Control and T2D groups increasing the accuracy to: 0.84. A total of 30 features, including the above-mentioned features selected by all ML models, were examined to find the minimal number of proteins that are necessary for correct classification of the patients into disease groups. We took the five proteins indicated by all ML models, plus GUSB as a core protein set, and added each protein one-by one and assessed the improvement in model performance. Four proteins (ADGRG2, IGFBP2, LPL, and PCOLCE), out of thirty, improved the AUC values resulting in a panel of 10 proteins. Using this panel of proteins, the accuracy increased to 0.96 with 0.967 macro-average sensitivity and 0.976 macro-average specificity (Figure 4, Supplementary Figure 2).

**Figure 4.**
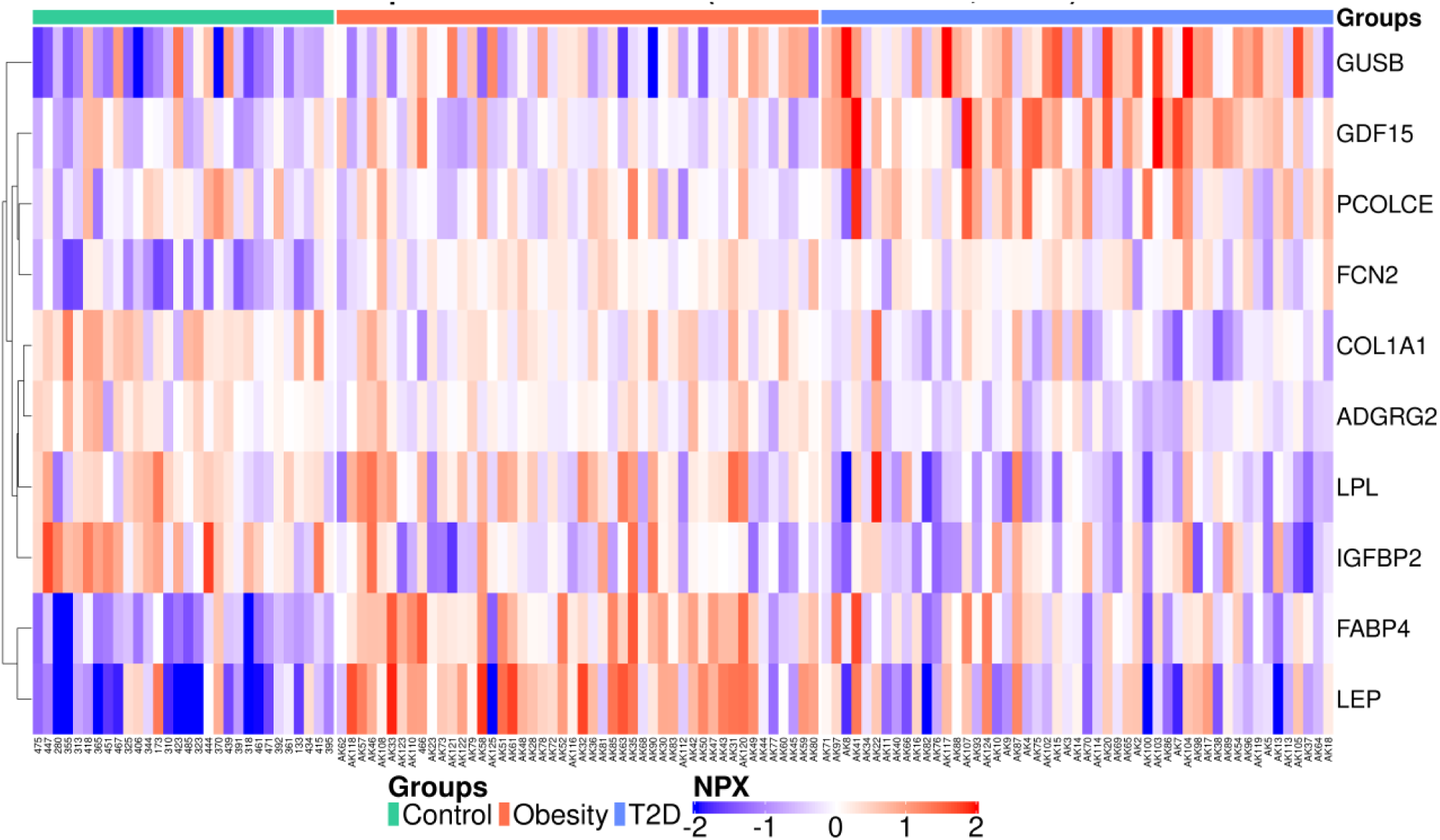
Heatmap of candidate biomarkers. Heatmaps show NPX values (blue=low expression, red=high expression) across samples in all clinical groups. The annotation bar above the heatmaps indicates clinical groups (green = Control, orange = Obesity, blue = T2D groups).

### 3.2. Verification of the candidate biomarkers on independent datasets

The panel of 10 potential biomarkers was verified on a Human Protein Atlas dataset comprising data from 834 patients grouped as Healthy Controls (n=834), Obesity (n=243), T2D (n=75) and Other (n=678). This later group contains samples from patients with MASLD, coronary artery calcification, and metabolic syndrome. The multiclass classification model demonstrated strong discriminative performance between the Healthy Control, Obesity and T2D groups, with a macro-average AUC of 0.857 and a micro-average AUC of 0.865 (Figure 5).

**Figure 5.**
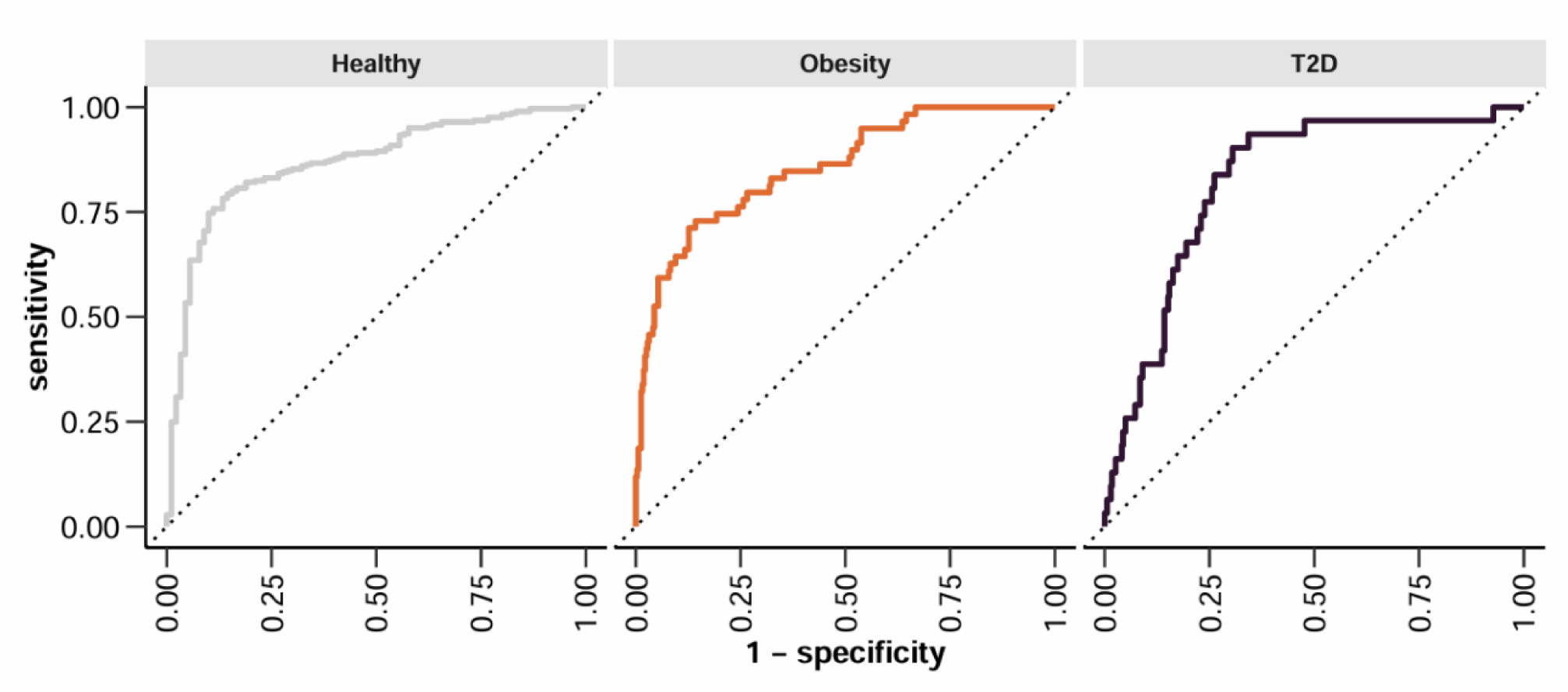
Receiver operating characteristic (ROC) curves for validation of the candidate biomarkers. Each panel shows the ROC curve for one class in a one-vs-rest multiclass classification framework. The diagonal dashed line indicates the performance of a random classifier.

The performance was a bit lower when besides the Obesity and T2D groups a composite of Healthy Control and Other groups was used. In this case the macro-average AUC was 0.743 and the micro-average AUC 0.724. Although the protein amounts between the studies are not directly comparable, the relative changes of proteins compared to Control in the HPA dataset was consistent with the changes observed in the original dataset (Figure 6).

**Figure 6.**
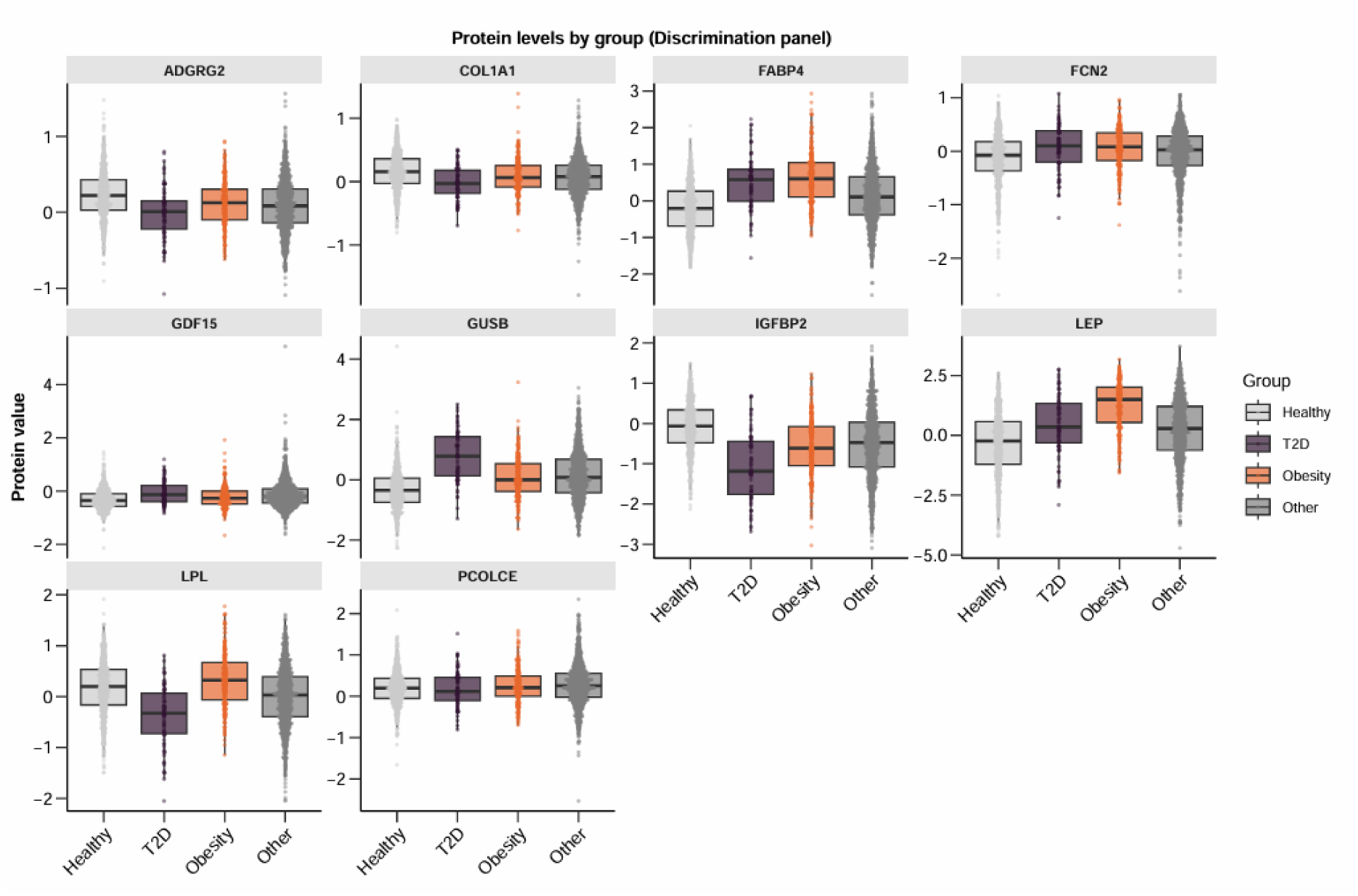
Distribution of candidate biomarker levels across groups in the HPA validation dataset. Boxplots showing the normalized protein abundance of the 10 candidate biomarkers across four groups in the HPA dataset: Healthy Controls, T2D, Obesity, and Other. Points represent individual samples, and boxplots indicate the median and IQR of protein amounts.

According to this, FCN2 and FABP4 can be considered as pan-disease marker with elevated levels in all disease groups. LPL and LEP can be considered as discriminatory proteins between the groups. The level of LPL decreased in T2D and increased in the Obesity group, while in the case of LEP elevation was observed in both groups with a marked increase in Obesity group. Decreased levels of PCOLCE in T2D group, and a decrease in all disease groups with a marked decrease in T2D was observed in case of ADGRG2, COL1A1, and IGFBP2. The GDF15 and GUSB levels were higher in all disease groups with further increase in T2D groups.

### 3.3. Unsupervised cluster analysis revealed molecular heterogeneity within the study groups

To explore potential molecular heterogeneity within the individual study groups, we performed unsupervised cluster analysis based on the proteomic profiles.

#### 3.3.1. Intragroup clusters show patient stratification

To examine the molecular heterogeneity of the study groups, recruited as homogenous from clinical point of view, k-means cluster analysis was applied. Our cluster analysis revealed distinct subgroups (clusters) within each study group, Control, Obesity, and T2D. The number of centroids was found to be optimal and set to *k =* 2 in each group (Figure 7, Supplementary Figure 3).

**Figure 7.**
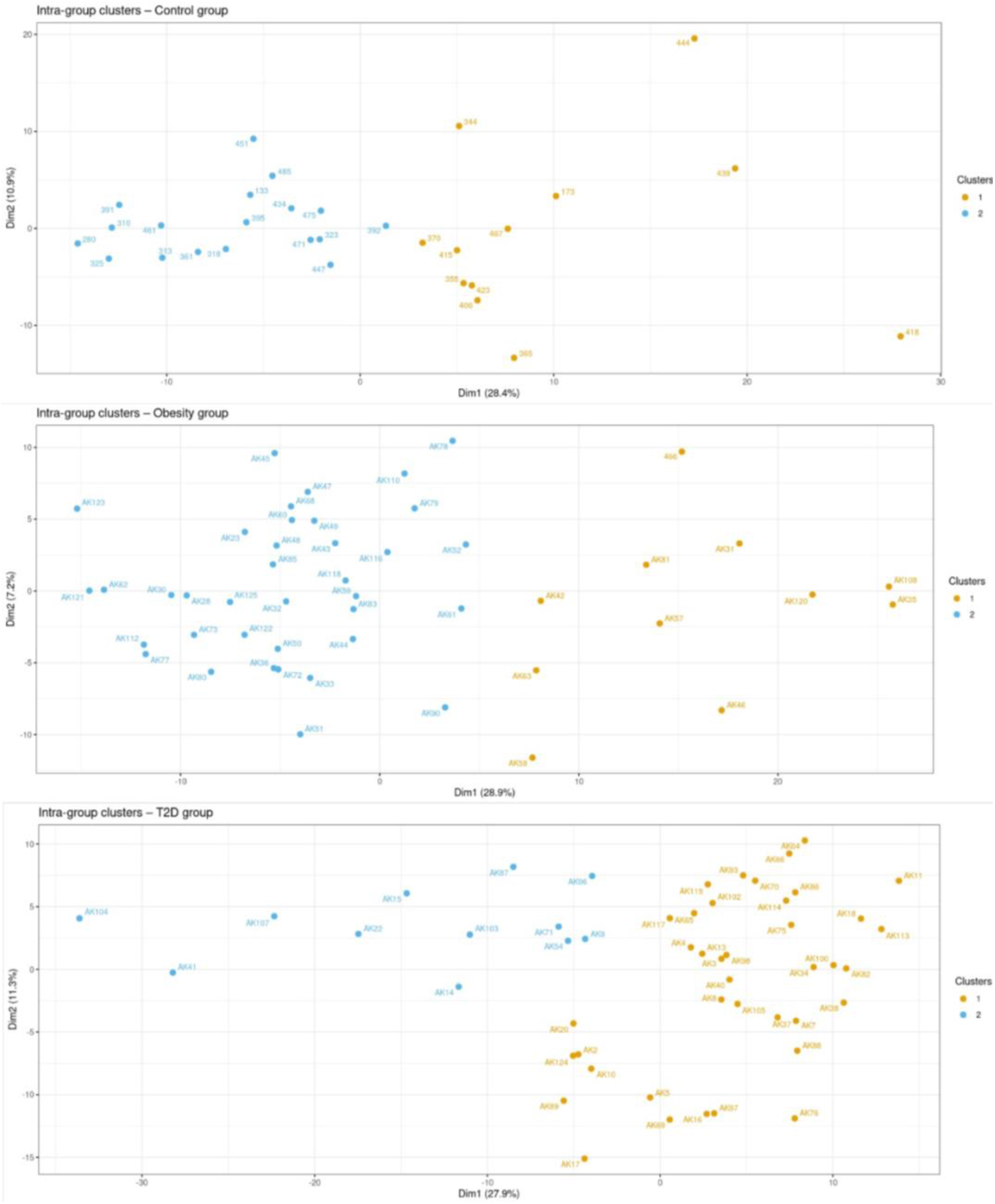
Two-dimensional visualization of intra-group clusters for three groups. Each point represents an individual, positioned according to the k-means clustering (*k* = 2) and colored by cluster assignment. The optimal number of clusters was determined by measuring average silhouette width at different number of clusters.

In the Control group, two clusters were identified, consisting of 18 and 12 individuals, respectively. The clusters were clearly separated along the 2D visualization, suggesting underlying heterogeneity within the Control population. In the Obesity group, clustering also produced two subgroups: one smaller cluster with 11 individuals and a larger one with 37 individuals. In case of the T2D group, two clusters of markedly different sizes were identified: a large cluster of 39 individuals and a smaller cluster of 12 individuals. Compared with individuals in Control group and in Obesity group, patients with T2D showed a stronger imbalance in cluster distribution, with most individuals concentrated in the cluster 1. The analysis performance was tested by the 5-fold CV method, and the results are listed in the Supplementary Table 3.

Together, these results show the presence of molecular heterogeneity across all clinical groups. While clustering yielded two subgroups in each group, the relative sizes of clusters varied substantially, with more balanced distributions in the Control group and imbalanced distributions in the Obesity and T2D groups. The separation between clusters was visibly distinct in the 2D plot graphs, and the calculated errors and accuracy were lower, corresponding to the distinction (Supplementary Table 3).

#### 3.3.2. Proteins responsible for subgroup discrimination

To further characterize the proteins driving the unsupervised k-means clusters, we applied the bootstrapped RF method^47–50^ to find the important proteins identifying the subgroups and listed them according to their MDGini scores. To establish objective thresholds for feature selection, we compared real data against null models derived from shuffled datasets, allowing us to distinguish biologically meaningful signals from noise. Model performance was evaluated across 1000 bootstrap iterations (Supplementary Table 3).

Mean OOB error rate of Control group was higher than of other two groups while the error was lowest in the T2D group. The mean class error per cluster, another indicator, was also lower in one of two clusters per group. Meanwhile, the average accuracy was between 0.955 and 0.982, and the AUC was approximately 1.00 in all groups, indicating that the RF model can be a model of choice to further examine these clusters.

The feature importance was measured by the MDGini score, averaged across bootstrap iterations. The higher the MDGini score the higher importance of the protein (Figure 8).

**Figure 8.**
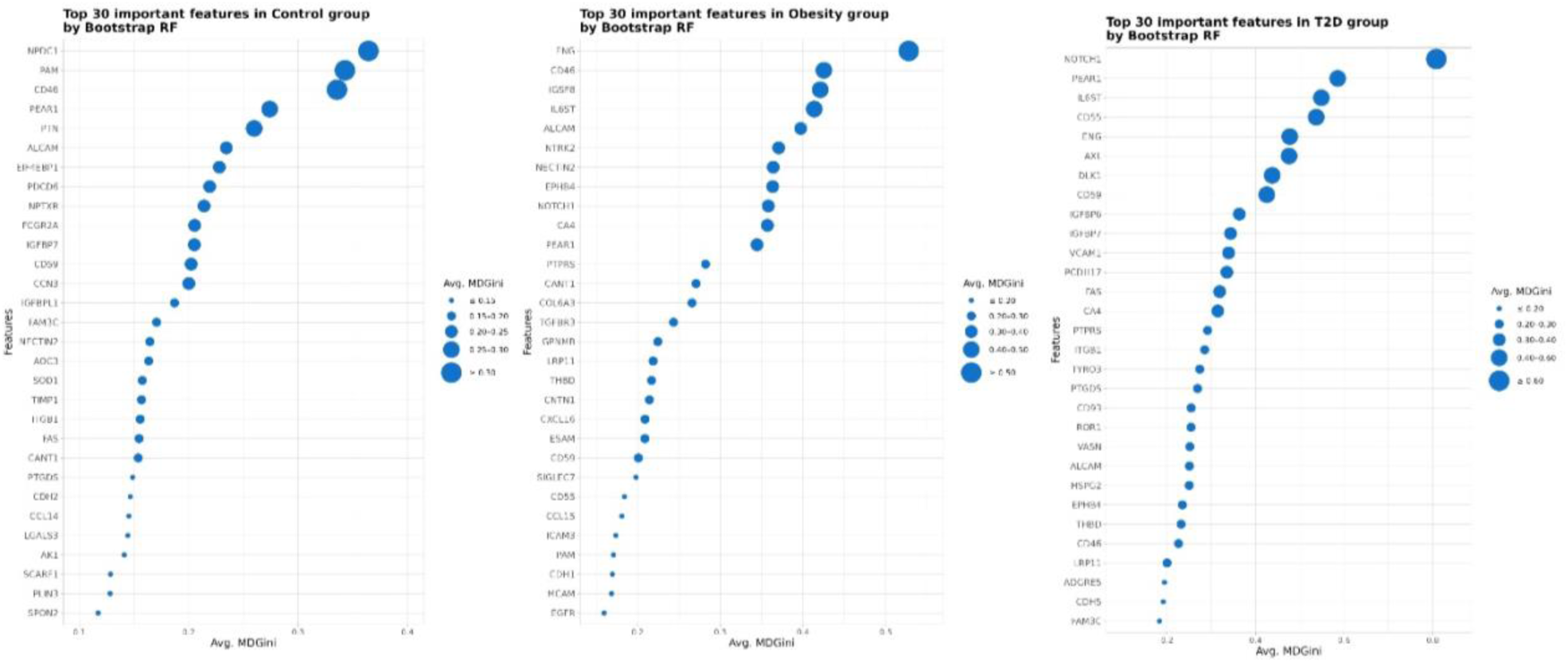
The top 30 important features for discriminating clusters within each group, as determined by bootstrap RF analysis. Feature importance was ordered by average MDGini across 1000 bootstrap iterations. The dot size corresponds to average MDGini intervals, with larger dots indicating higher importance.

Using the data-driven threshold, we identified the majority of proteins as being more informative than expected by chance. Specifically, 311/366 proteins in the Control group, 251/366 proteins in the Obesity group, and 305/366 proteins in the T2D group were above this threshold. This indicates that a large proportion of the proteins contributes at least modestly to the discrimination of clusters within each group.

The features with high importance can be unstable and show up in a few bootstrap samples. Therefore, we tested if those top 30 most important features are repeatedly selected as top important features in at least 70% percent of the bootstrap samples. If they were selected as top important features more than 700 times out of 1000 bootstraps, they were considered highly stable important features (Figure 9).

**Figure 9.**
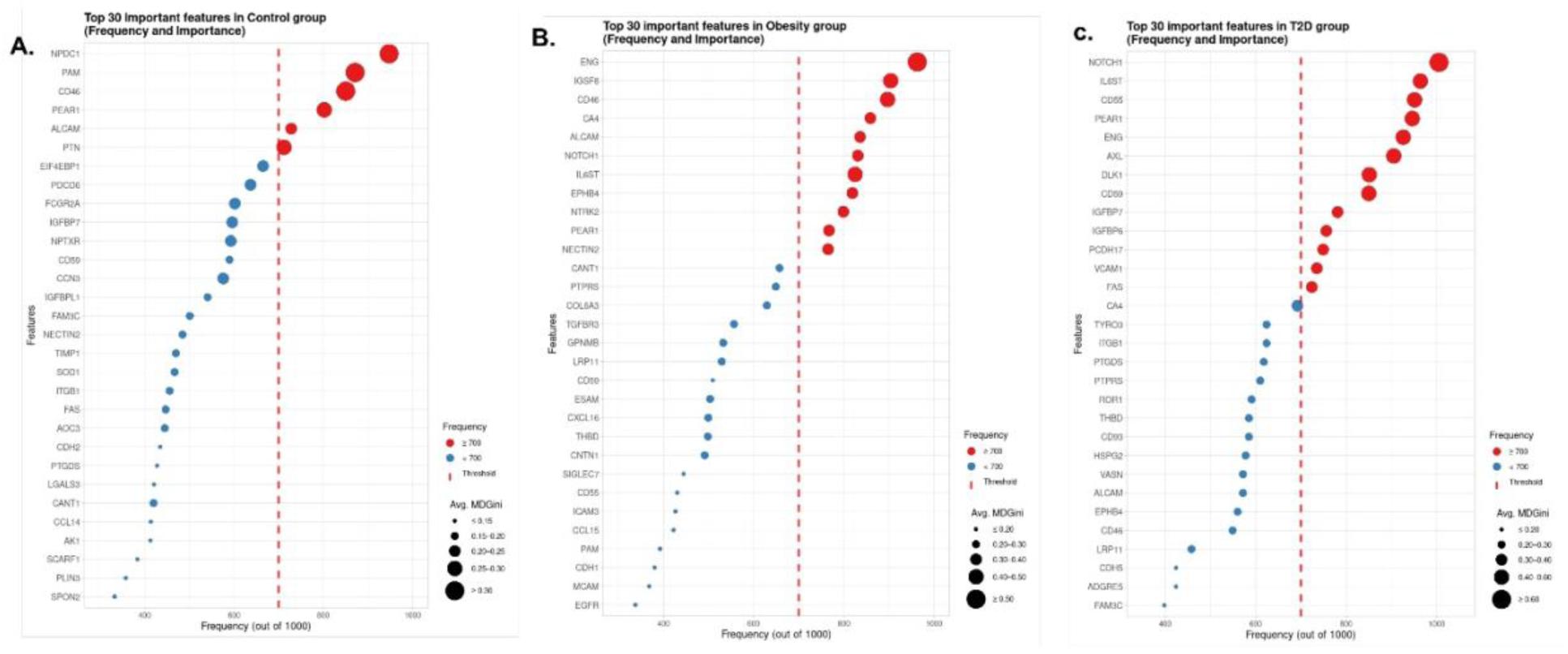
Frequency and importance of selected featured in each group (A-C). The scaled dot plot shows the top important features ranked by average MDGini score across 1000 bootstrap RF iterations. The x-axis represents the frequency of each protein appeared at y-axis, and the dot size corresponds to the average MDGini score ranges. Proteins at the right side of the red dashed line are considered highly stable important features, colored in red, while red dashed line indicates the stability threshold (frequency ≥ 700/1000).

In total, seventeen proteins were identified as highly stable important features in all three groups and consistently distinguished two clusters in each group. One protein, PEAR1, was common in all three groups, two proteins, ALCAM and CD46 were shared between Control and Obesity groups, three proteins, ENG, IL6ST, NOTCH1, were common in the Obesity and T2D groups. There was no shared protein between the Control and T2D groups.

Out of six, three proteins, NPDC1, PAM, and PTN, were unique in Control group. Eleven proteins were highly stable in Obesity group and five of them, CA4, EPHB4, IGSF8, NECTIN2, and NTRK2 were unique in this group. In T2D group, thirteen proteins were identified as highly stable important features and nine of them, AXL, CD55, CD59, DLK1, FAS, IGFBP6, IGFBP7, PCDH17, and VCAM1 were unique in this group (Supplementary Figure 4).

The heatmaps visualize the expression pattern of these highly stable important features across individuals, stratified by intragroup clusters (Figure 10, A-C). All those features are displaying a clear separation between the two clusters in each group. Corresponding boxplots also show the significant differences in protein expression between clusters for all identified features (Figure 10, D-F). These results demonstrate that intragroup clustering is supported by a small but robust set of highly stable important features, with increasing molecular complexity observed in our study groups. The consistency between heatmap patterns and boxplot-based statistical comparisons underscores the reliability of these proteins as intragroup discriminative markers.

**Figure 10.**
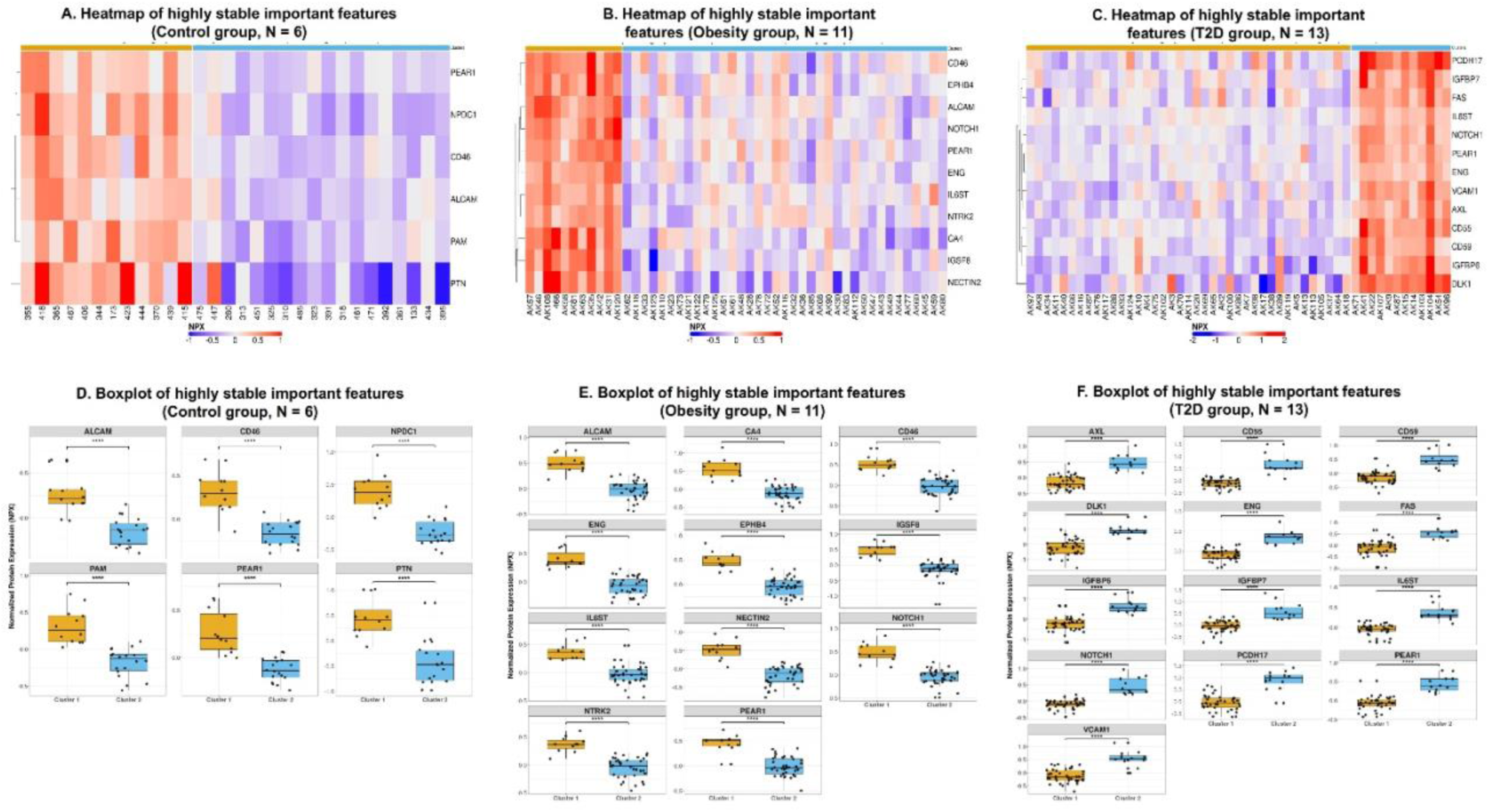
Proteins that drive subgroup discrimination within each study group .Proteins identified as highly stable important features in Control group (A and D), in Obesity group (B and E), in T2D group (C and F). (A-C) Heatmaps displaying NPX values of the highly stable important features identified by bootstrap RF. The heatmap columns represent individual participants, separated into two clusters based on the k-means cluster assignment (*k* = 2), and rows represent proteins. Color intensity corresponds to NPX levels, with red indicating higher and blue indicating lower levels. (D-F) Boxplots showing the NPX values of proteins in clusters. y-axis represents the NPX values and the x-axis represents the clusters with color code, **** indicates adj.p-value < 0.0001. Clusters are indicated by the colors, orange color represents cluster 1 and blue color represents cluster 2.

#### 3.3.3. The functional examination of proteins reflecting molecular heterogeneity in the study groups

To investigate the biological function of highly stable proteins discriminating between the subgroups, GO biological process enrichment analysis was performed separately within each group. The enriched GO terms were visualized using a Sankey diagram, illustrating the relationships between each group and the biological processes associated with their highly stable important features (Supplementary Figure 5). In the Control group, enriched processes were primarily related to the regulation of hematopoiesis and cell development. The proteins reflecting the heterogeneity in the Obesity group were mostly related to cell development, epithelial to mesenchymal transition, extracellular matrix remodeling, vasculogenesis and NOTCH signaling. At the same time, the proteins responsible for cluster discrimination in T2D group had a role in hemostasis and immune functions, some of which can be linked to blood coagulation. Due to overlapping proteins, some of the functions observed in the Obesity group appeared in T2D group as well.

It is well known that in Obesity the prevalence of MASLD is much higher^51,52^, we examined the role of individual proteins in liver disease. Many of them were already identified in conjunction to liver disease^53–55^. To test if these subgroups are associated with the liver condition, we calculated LiverRisk Score^14^ using the internet-based tool (https://www.liverriskscore.com/#/home) (Figure 11).

**Figure 11.**
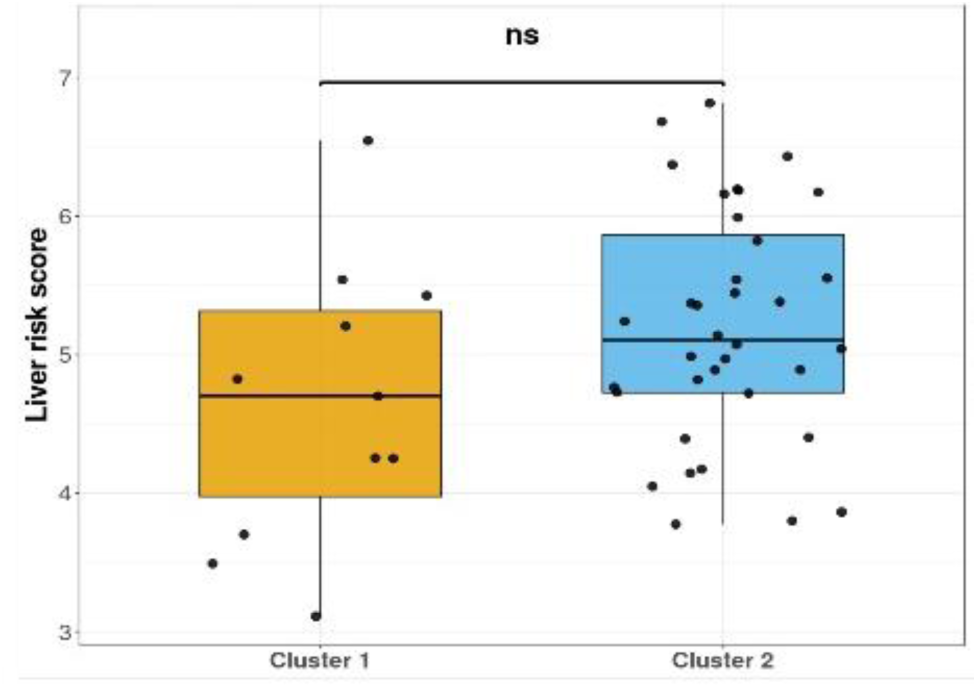
LiverRisk Score calculated in the case of patients belonging to the two clusters within the Obesity group. y-axis represents the LiverRisk Score and the x-axis represents the clusters with color code. Clusters are indicated by colors; orange color represents cluster 1 and blue color represents cluster 2.

Although an increasing trend was observed in case of LiverRisk Scores in cluster 2, compared to cluster 1, there was no statistically significant difference in the LiverRisk Scores between the subgroups (p-value=0.099).

#### 3.3.4. The disease groups are homogenous based on their clinical profile

In addition to the cluster analysis, we integrated proteomics-driven clusters with clinical variables to assess whether our identified intragroup clusters correspond to distinct clinical phenotypes. We compared two clusters within each group using routinely measured clinical laboratory variables. The comparisons of the available clinical parameters did not reveal statistically significant differences between the identified clusters (Supplementary Figure 6), suggesting that proteomics-based ML-driven heterogeneity was not captured by routine clinical measurements in this study.

## 4. Discussion

Obesity is associated to chronic, low-grade inflammation, often called meta-inflammation^7^ and over time it is transitioned to T2D if multiple factors emerge^56–62^. We applied several supervised ML models to reveal the molecular distinction between the Control, Obesity, and T2D groups and identified candidate biomarkers able to distinguish the clinical groups from each other. ML models are widely used in recent proteomics-based studies for metabolic disorders^57–60^ in order to reveal the signature proteins and predict the early onset of diseases.

Across four different ML models (RF, LASSO, SVM-RFE, and VE), five proteins: FABP4, LEP, GDF15, FCN2, and COL1A1, were consistently selected as important discriminatory features. The algorithmically distinct methods (tree-based, linear-regularized, margin-based, and ensemble SHAP-based) strongly support the robustness of those candidate biomarkers. In addition to them, we expanded our list with ADGRG2, GUSB, IGFBP2, LPL, and PCOLCE as they increased the discrimination efficiency between the groups.

The candidate biomarker panel identified in our study contains proteins that were previously identified either as biomarkers for Obesity and/or T2D or a strong correlation between their amount and the presence of the pathological condition was demonstrated. According to the DIETFITS and EpiHealth studies, LEP, FABP4, GDF-15, IGFBP-2, and LPL were among the positively associated proteins with BMI in patients with Obesity or overweight at baseline, as well as with changes in BMI during 6 months with LEP and FABP4 showing the strongest associations^61^. FABP4 and LEP are well-known adipokines, the master regulators of adipogenesis and insulin responsiveness^62^. Elevated circulating FABP4 and LEP levels have been repeatedly associated with Obesity and T2D^63–66^. The EpiHealth study revealed that the elevated levels of FABP4 and GDF15 were associated with an increased risk of T2D, whereas higher levels of LPL and IGFBP2 were associated with lower risk of T2D^67^, suggesting that LPL and IGFBP2 may serve as protective factors. Another study revealed that IGFBP2 expression is regulated by LEP and can serve as a therapeutic target in T2D^68^. GDF15 and IGFBP2 were among those proteins associated with T2D^69^ which corroborates with our ML model-identified results. Consistent selection of these proteins by all models aligns with their role as mediators between adipose tissue inflammation and systemic metabolic alterations leading to T2D^70,71^.

Chronic metabolic stress is known to induce low-grade fibrosis and endothelial dysfunction, both hallmarks of T2D progression^72^. COL1A1 and PCOLCE, along with additional vascular-associated proteins identified in model-specific selections, may be indicators of extracellular matrix remodeling and vascular adaptation^73,74^. Down-regulation of COL1A1 by natural ingredients used for hepatic fibrosis treatment *in vivo*, suggested that they contribute to chronic liver diseases and named as fibrogenic markers^75,76^. It was earlier identified that Obesity and T2D both increase the risk of MASLD (formerly, Non-Alcoholic Fatty Liver Disease, NAFLD)^77^. By the *in silico* funnel strategy, FCN2 was, either alone or combined with other parameters, identified as a candidate biomarker of fibrosis, and its plasma level was inversely correlated with the stage of liver fibrosis in subjects with Obesity^78^.

The potential biomarker panel was tested on an independent dataset containing information acquired with Olink Explore 1536 panel on donors recruited to SCAPIS and IGTM studies (HPA dataset). As expected, the AUC values higher than 0.9, very likely generated by overfitting of the data due to relatively low sample size in our original dataset, could not be reproduced, but AUC values of 0.85-0.86 show the good discriminatory power of the candidate biomarker panel on a different dataset.

Our panel, verified on larger dataset originating from independent studies demonstrates its potential in discriminating the clinical groups from each other.

Another clinically relevant question is how heterogeneous a patient group is. For a more personalized and targeted therapy, information on the potential subgroups is critically important. The groups recruited into this study were regular patient groups, and extra care was taken to generate groups that are homogenous from clinical point of view. This homogeneity could be confirmed by our analyses, where no statistically significant difference in clinical variables could be observed within groups. To examine the molecular heterogeneity, unsupervised k-means clustering was applied and identified two clusters within each clinical group based on the examined proteins. The high bootstrap stability and low OOB error rates confirmed that these clusters are not random partitions but reflect reproducible proteomic signatures. This indicates that routine laboratory and anthropometric measures may lack sensitivity to detect underlying molecular subtypes of these conditions, highlighting the added value of proteomics-driven stratification.

The Control group showed balanced clusters enriched for proteins having a role in hematopoietic and cell development pathways. The clusters in the Obesity group could be discriminated from each other based on proteins that demonstrated enriched functions such as vasculogenesis and NOTCH-related pathways whereas, T2D subgroups were discriminated based on proteins having a role in blood coagulation and complement activation. Our unsupervised cluster analysis revealed that eleven proteins were highly stable to discriminate the two clusters in the Obesity group. Among those proteins, EPHB4 and CD166 antigen (ALCAM) were found to be positively associated with high BMI and their blood level decreased after patients with Obesity or overweight lost weight according to the cross-sectional analysis by DIETFITS and EpiHealth cohorts^61^. As the prevalence of liver disease is higher in Obesity^79,80^, each protein was checked if they were previously associated with liver cirrhosis or MASLD. For instance, ENG plays an important role in liver fibrosis by altering the TGF-β signaling via the ALK1-Smad1/5 and ALK-Smad2/3 pathways^81^ and was identified as a non-invasive biomarker for the MASLD^82^. ENG, NTRK2, and AXL were differentially upregulated at the late stage of progressive fibrosis in MASLD, while IGSF8 was among those in the early stage^83^. Calculating the LiverRisk Scores, the two subgroups showed a clear, but statistically not significant distinction, and the overall values for the whole Obesity group indicated a low or moderate risk for liver disease^14^.

Some of the proteins, such as ENG, IL6ST, NOTCH1, and PEAR1 were common between the protein sets, enabling clustering both in the Obesity and T2D groups, suggesting the presence of multiple possible complications in both groups.

Regarding the proteins discriminating the subgroups in T2D, many of them were associated with T2D-related complications, especially vasculature-related complications. VCAM1 is a marker of microvascular complications acting as a key mediator of endothelial dysfunction and inflammation^84^. CD55 and CD59 both regulate complement activation; the level of glycated CD59 is elevated in patients with microvascular complications^85^, and the expression of both CD55 and CD59 was found to be significantly lower in patients with nephropathy, retinopathy, and cardiovascular disease compared to Controls^86^. Multiple studies demonstrated the link between the IGFBP7 and the complications of T2D^87^. Examining the data from the BIOSTAT-CHF cohort, IGFBP7 was identified as a marker of cardiovascular complications; its levels strongly correlated with the presence of atrial fibrillation, T2D and worse clinical status^88^. Regarding IGFBP6, its level was correlated to complications in Type 1 Diabetes^89^, but not T2D. Elevated levels of AXL were shown to be linked to impaired vascularization, leading potentially to microvasculature complications and retinopathy^90^. The association of these proteins with the more severe, especially vasculature-related complications, pinpoints the importance of definition of subgroups in the T2D group considered as homogenous from a clinical point of view. Patient stratification can help clinicians to define subgroups with different therapeutic needs allowing the application of more personalized treatment protocols.

Our findings support the importance of proteomics analyses beside of the classical clinical variables to depict the molecular heterogeneity present inside the groups considered as homogenous from a clinical point of view.

## 5. Limitations

The relatively small sample size in our high-dimensional proteomics data introduces a risk of overfitting and limits model generalizability. Although multiple resampling strategies including bootstrap aggregation, OOB error estimation, and repeated k-fold cross-validation were implemented, these approaches primarily provide internal validation and may not fully capture performance in independent datasets.

To reduce the false associations, feature importance was benchmarked against null distributions derived from label-permuted data, and only features demonstrating consistent importance and stability across bootstrap iterations were retained. Nonetheless, high-dimensional proteomic data remain susceptible to instability in feature selection, particularly in the presence of correlated protein expression patterns and potential technical variability.

## 6. Conclusion

Obesity and T2D are heterogeneous metabolic disorders whose molecular diversity is incompletely captured by conventional clinical parameters. To address this gap, we applied complementary supervised and unsupervised ML approaches to high-dimensional circulating protein profiles to investigate molecular heterogeneity across Control, Obesity, and T2D groups. Supervised models identified protein signatures that robustly discriminated between clinical groups and highlighted candidate biomarkers, verified on Human Protein Atlas dataset. Unsupervised clustering further revealed distinct molecular subgroups within each diagnostic category, indicating substantial group-level molecular heterogeneity not reflected by standard clinical variables. Notably, subgroup analysis within the Obesity group identified proteomic patterns associated with increased risk of liver fibrosis, while within the T2D group, with altered blood coagulation. These findings demonstrate that integrating ML with proteomics can capture the molecular heterogeneity in Obesity and T2D groups and may enable improved patient stratification and biomarker discovery for metabolic complications.

## Acknowledgements

We are very thankful for Dr. Endre Kristóf for reviewing our manuscript and for his critical comments. We also thank Uladzislau Vododokhau for his initial script drafting.

## References

(1) IDF Diabetes Atlas, 10th edition.; International Diabetes Federation: Brussels, 2021.

(2) Dai, W.; Jiang, L. Dysregulated Mitochondrial Dynamics and Metabolism in Obesity, Diabetes, and Cancer. Front. Endocrinol. 2019, 10, 570. 10.3389/fendo.2019.00570.

(3) Lee, M.-J.; Wu, Y.; Fried, S. K. Adipose Tissue Heterogeneity: Implication of Depot Differences in Adipose Tissue for Obesity Complications. Mol. Aspects Med. 2013, 34 (1), 1–11. 10.1016/j.mam.2012.10.001.

(4) Nagarajan, S. R.; Cross, E.; Sanna, F.; Hodson, L. Dysregulation of Hepatic Metabolism with Obesity: Factors Influencing Glucose and Lipid Metabolism. Proc. Nutr. Soc. 2022, 81 (1), 1–11. 10.1017/S0029665121003761.

(5) Esser, N.; Legrand-Poels, S.; Piette, J.; Scheen, A. J.; Paquot, N. Inflammation as a Link between Obesity, Metabolic Syndrome and Type 2 Diabetes. Diabetes Res. Clin. Pract. 2014, 105 (2), 141–150. 10.1016/j.diabres.2014.04.006.

(6) Rohm, T. V.; Meier, D. T.; Olefsky, J. M.; Donath, M. Y. Inflammation in Obesity, Diabetes, and Related Disorders. Immunity 2022, 55 (1), 31–55. 10.1016/j.immuni.2021.12.013.

(7) Pereira, S. S.; Alvarez-Leite, J. I. Low-Grade Inflammation, Obesity, and Diabetes. Curr. Obes. Rep. 2014, 3 (4), 422–431. 10.1007/s13679-014-0124-9.

(8) Piché, M.-E.; Tchernof, A.; Després, J.-P. Obesity Phenotypes, Diabetes, and Cardiovascular Diseases. Circ. Res. 2020, 126 (11), 1477–1500. 10.1161/CIRCRESAHA.120.316101.

(9) Ulambayar, B.; Ghanem, A. S.; Kovács, N.; Trefán, L.; Móré, M.; Nagy, A. C. Cardiovascular Disease and Risk Factors in Adults with Diabetes Mellitus in Hungary: A Population-Based Study. Front. Endocrinol. 2023, 14, 1263365. 10.3389/fendo.2023.1263365.

(10) Lundberg, M.; Eriksson, A.; Tran, B.; Assarsson, E.; Fredriksson, S. Homogeneous Antibody-Based Proximity Extension Assays Provide Sensitive and Specific Detection of Low-Abundant Proteins in Human Blood. Nucleic Acids Res. 2011, 39 (15), e102–e102. 10.1093/nar/gkr424.

(11) Bertalan, P. M.; Nokhoijav, E.; Pap, Á.; Neagu, G. C.; Káplár, M.; Darula, Z.; Kalló, G.; Prokai, L.; Csősz, É. Comprehensive Insights into Obesity and Type 2 Diabetes from Protein Network, Canonical Pathway, Phosphorylation and Antimicrobial Peptide Signatures of Human Serum. Proteomes 2025, 13 (4), 67. 10.3390/proteomes13040067.

(12) Vadadokhau, U.; Varga, I.; Káplár, M.; Emri, M.; Csősz, É. Examination of the Complex Molecular Landscape in Obesity and Type 2 Diabetes. Int. J. Mol. Sci. 2024, 25 (9), 4781. 10.3390/ijms25094781.

(13) Nokhoijav, E.; Guba, A.; Kumar, A.; Kunkli, B.; Kalló, G.; Káplár, M.; Somodi, S.; Garai, I.; Csutak, A.; Tóth, N.; Emri, M.; Tőzsér, J.; Csősz, É. Metabolomic Analysis of Serum and Tear Samples from Patients with Obesity and Type 2 Diabetes Mellitus. Int. J. Mol. Sci. 2022, 23 (9), 4534. 10.3390/ijms23094534.

(14) Liu, S.; Chen, X.; Jiang, X.; Yin, X.; Fekadu, G.; Liu, C.; He, Y.; Chen, H.; Ni, W.; Wang, R.; Zeng, Q.-L.; Chen, Y.; Yang, L.; Shi, R.; Ju, S.-H.; Shen, J.; Gao, J.; Zhao, L.; Ming, W.; Zhong, V. W.; Teng, G.-J.; Qi, X. LiverRisk Score: An Accurate, Cost-Effective Tool to Predict Fibrosis, Liver-Related, and Diabetes-Related Mortality in the General Population. Med 2024, 5 (6), 570–582.e4. 10.1016/j.medj.2024.03.003.

(15) Wik, L.; Nordberg, N.; Broberg, J.; Björkesten, J.; Assarsson, E.; Henriksson, S.; Grundberg, I.; Pettersson, E.; Westerberg, C.; Liljeroth, E.; Falck, A.; Lundberg, M. Proximity Extension Assay in Combination with Next-Generation Sequencing for High-Throughput Proteome-Wide Analysis. Mol. Cell. Proteomics 2021, 20, 100168. 10.1016/j.mcpro.2021.100168.

(16) Breiman, L. Random Forests. Mach. Learn. 2001, 45 (1), 5–32. 10.1023/A:1010933404324.

(17) Breiman, L.; Cutler, A.; Liaw, A.; Wiener, M. randomForest: Breiman and Cutlers Random Forests for Classification and Regression, 2002, 4.7–1.2. 10.32614/CRAN.package.randomForest.

(18) Tibshirani, R. Regression Shrinkage and Selection Via the Lasso. J. R. Stat. Soc. Ser. B Stat. Methodol. 1996, 58 (1), 267–288. 10.1111/j.2517-6161.1996.tb02080.x.

(19) Tay, J. K.; Narasimhan, B.; Hastie, T. Elastic Net Regularization Paths for All Generalized Linear Models. J. Stat. Softw. 2023, 106 (1). 10.18637/jss.v106.i01.

(20) Friedman, J.; Hastie, T.; Tibshirani, R. Regularization Paths for Generalized Linear Models via Coordinate Descent. J. Stat. Softw. 2010, 33 (1). 10.18637/jss.v033.i01.

(21) Cortes, C.; Vapnik, V. Support-Vector Networks. Mach. Learn. 1995, 20 (3), 273–297. 10.1007/BF00994018.

(22) Karatzoglou, A.; Meyer, D.; Hornik, K. Support Vector Machines in *R*. J. Stat. Softw. 2006, 15 (9). 10.18637/jss.v015.i09.

(23) Meyer, D.; Dimitriadou, E.; Hornik, K.; Weingessel, A.; Leisch, F. E1071: Misc Functions of the Department of Statistics, Probability Theory Group (Formerly: E1071), TU Wien, 1999, 1.7–16. 10.32614/CRAN.package.e1071.

(24) Lundberg, S. M.; Lee, S.-I. A Unified Approach to Interpreting Model Predictions. 2017, 4765–4774.

(25) Sesmero, M. P.; Ledezma, A. I.; Sanchis, A. Generating Ensembles of Heterogeneous Classifiers Using Stacked Generalization. WIREs Data Min. Knowl. Discov. 2015, 5 (1), 21–34. 10.1002/widm.1143.

(26) Uhlén, M.; Fagerberg, L.; Hallström, B. M.; Lindskog, C.; Oksvold, P.; Mardinoglu, A.; Sivertsson, Å.; Kampf, C.; Sjöstedt, E.; Asplund, A.; Olsson, I.; Edlund, K.; Lundberg, E.; Navani, S.; Szigyarto, C. A.-K.; Odeberg, J.; Djureinovic, D.; Takanen, J. O.; Hober, S.; Alm, T.; Edqvist, P.-H.; Berling, H.; Tegel, H.; Mulder, J.; Rockberg, J.; Nilsson, P.; Schwenk, J. M.; Hamsten, M.; Von Feilitzen, K.; Forsberg, M.; Persson, L.; Johansson, F.; Zwahlen, M.; Von Heijne, G.; Nielsen, J.; Pontén, F. Tissue-Based Map of the Human Proteome. Science 2015, 347 (6220), 1260419. 10.1126/science.1260419.

(27) Wu, H.; Tremaroli, V.; Schmidt, C.; Lundqvist, A.; Olsson, L. M.; Krämer, M.; Gummesson, A.; Perkins, R.; Bergström, G.; Bäckhed, F. The Gut Microbiota in Prediabetes and Diabetes: A Population-Based Cross-Sectional Study. Cell Metab. 2020, 32 (3), 379–390.e3. 10.1016/j.cmet.2020.06.011.

(28) Bergström, G.; Berglund, G.; Blomberg, A.; Brandberg, J.; Engström, G.; Engvall, J.; Eriksson, M.; De Faire, U.; Flinck, A.; Hansson, M. G.; Hedblad, B.; Hjelmgren, O.; Janson, C.; Jernberg, T.; Johnsson, Å.; Johansson, L.; Lind, L.; Löfdahl, C. -G.; Melander, O.; Östgren, C. J.; Persson, A.; Persson, M.; Sandström, A.; Schmidt, C.; Söderberg, S.; Sundström, J.; Toren, K.; Waldenström, A.; Wedel, H.; Vikgren, J.; Fagerberg, B.; Rosengren, A. The Swedish CArdioPulmonary BioImage Study: Objectives and Design. J. Intern. Med. 2015, 278 (6), 645–659. 10.1111/joim.12384.

(29) Ikotun, A. M.; Ezugwu, A. E.; Abualigah, L.; Abuhaija, B.; Heming, J. K-Means Clustering Algorithms: A Comprehensive Review, Variants Analysis, and Advances in the Era of Big Data. Inf. Sci. 2023, 622, 178–210. 10.1016/j.ins.2022.11.139.

(30) Steinley, D.; Brusco, M. J. Initializing K-Means Batch Clustering: A Critical Evaluation of Several Techniques. J. Classif. 2007, 24 (1), 99–121. 10.1007/s00357-007-0003-0.

(31) Hartigan, J. A.; Wong, M. A. Algorithm AS 136: A K-Means Clustering Algorithm. Appl. Stat. 1979, 28 (1), 100. 10.2307/2346830.

(32) Lassané, L. KMEANS.KNN: KMeans and KNN Clustering Package, 2024, 0.1.0. 10.32614/CRAN.package.KMEANS.KNN.

(33) Henderson, A. R. The Bootstrap: A Technique for Data-Driven Statistics. Using Computer-Intensive Analyses to Explore Experimental Data. Clin. Chim. Acta 2005, 359 (1–2), 1–26. 10.1016/j.cccn.2005.04.002.

(34) Calle, M. L.; Urrea, V. Letter to the Editor: Stability of Random Forest Importance Measures. Brief. Bioinform. 2011, 12 (1), 86–89. 10.1093/bib/bbq011.

(35) Kassambara, A.; Mundt, F. Factoextra: Extract and Visualize the Results of Multivariate Data Analyses, 2016, 1.0.7. 10.32614/CRAN.package.factoextra.

(36) Hadley Wickham. Ggplot2: Elegant Graphics for Data Analysis; Springer-Verlag New York, 2016. 10.1007/978-3-319-24277-4.

(37) Gu, Z. Complex Heatmap Visualization. iMeta 2022, 1 (3), e43. 10.1002/imt2.43.

(38) Xu, S.; Hu, E.; Cai, Y.; Xie, Z.; Luo, X.; Zhan, L.; Tang, W.; Wang, Q.; Liu, B.; Wang, R.; Xie, W.; Wu, T.; Xie, L.; Yu, G. Using clusterProfiler to Characterize Multiomics Data. Nat. Protoc. 2024, 19 (11), 3292–3320. 10.1038/s41596-024-01020-z.

(39) Yu, G.; Wang, L.-G.; Han, Y.; He, Q.-Y. clusterProfiler: An R Package for Comparing Biological Themes Among Gene Clusters. OMICS J. Integr. Biol. 2012, 16 (5), 284–287. 10.1089/omi.2011.0118.

(40) Marc Carlson. Org.Hs.Eg.Db: Genome Wide Annotation for Human, 2024. 10.18129/B9.bioc.org.Hs.eg.db.

(41) Haynes, W. Benjamini–Hochberg Method. In Encyclopedia of Systems Biology; Dubitzky, W., Wolkenhauer, O., Cho, K.-H., Yokota, H., Eds.; Springer New York: New York, NY, 2013; pp 78–78. 10.1007/978-1-4419-9863-7_1215.

(42) R Core Team. R: A Language and Environment for Statistical Computing, 2025. https://www.R-project.org/.

(43) Haynes, W. Wilcoxon Rank Sum Test. In Encyclopedia of Systems Biology; Dubitzky, W., Wolkenhauer, O., Cho, K.-H., Yokota, H., Eds.; Springer New York: New York, NY, 2013; pp 2354–2355. 10.1007/978-1-4419-9863-7_1185.

(44) Kruskal, W. H.; Wallis, W. A. Use of Ranks in One-Criterion Variance Analysis. J. Am. Stat. Assoc. 1952, 47 (260), 583–621. 10.1080/01621459.1952.10483441.

(45) McHugh, M. L. The Chi-Square Test of Independence. *Biochem*. Medica 2013, 143–149. 10.11613/BM.2013.018.

(46) Wickham, H.; Averick, M.; Bryan, J.; Chang, W.; McGowan, L.; François, R.; Grolemund, G.; Hayes, A.; Henry, L.; Hester, J.; Kuhn, M.; Pedersen, T.; Miller, E.; Bache, S.; Müller, K.; Ooms, J.; Robinson, D.; Seidel, D.; Spinu, V.; Takahashi, K.; Vaughan, D.; Wilke, C.; Woo, K.; Yutani, H. Welcome to the Tidyverse. J. Open Source Softw. 2019, 4 (43), 1686. 10.21105/joss.01686.

(47) Henkens, M. T. H. M.; Van Ommen, A.; Remmelzwaal, S.; Valstar, G. B.; Wang, P.; Verdonschot, J. A. J.; Hazebroek, M. R.; Hofstra, L.; Van Empel, V. P. M.; Beulens, J. W. J.; Den Ruijter, H. M.; Heymans, S. R. B. The HFA-PEFF Score Identifies ‘early-HFpEF’ Phenogroups Associated with Distinct Biomarker Profiles. ESC Heart Fail. 2022, 9 (3), 2032–2036. 10.1002/ehf2.13861.

(48) Wu, Y.; Liu, C.; Huang, J.; Wang, F. Quantitative Proteomics Reveals Pregnancy Prognosis Signature of Polycystic Ovary Syndrome Women Based on Machine Learning. Gynecol. Endocrinol. 2024, 40 (1), 2328613. 10.1080/09513590.2024.2328613.

(49) Misra, S.; Kawamura, Y.; Singh, P.; Sengupta, S.; Nath, M.; Rahman, Z.; Kumar, P.; Kumar, A.; Aggarwal, P.; Srivastava, A. K.; Pandit, A. K.; Mohania, D.; Prasad, K.; Mishra, N. K.; Vibha, D. Prognostic Biomarkers of Intracerebral Hemorrhage Identified Using Targeted Proteomics and Machine Learning Algorithms. PLOS ONE 2024, 19 (6), e0296616. 10.1371/journal.pone.0296616.

(50) Yasar, S.; Melekoglu, R. Proteomic Alterations in Ovarian Cancer—Predicting Residual Disease Status Using Artificial Intelligence and SHAP-Based Biomarker Interpretation. Front. Med. 2025, 12, 1562558. 10.3389/fmed.2025.1562558.

(51) Zhou, X.-D.; Targher, G.; Byrne, C. D.; Shapiro, M. D.; Chen, L.-L.; Zheng, M.-H. Metabolic Dysfunction-Associated Fatty Liver Disease: Bridging Cardiology and Hepatology. Cardiol. Plus 2024, 9 (4), 275–282. 10.1097/CP9.0000000000000106.

(52) Mandorfer, M.; Semmler, G.; Aigner, E.; Bräuer, A.; Maria Brix, J.; Clodi, M.; Datz, C.; Effenberger, M.; Felsenreich, D. M.; Ludvik, B.; Maieron, A.; Peck-Radosavljevic, M.; Ress, C.; Scherzer, T.-M.; Sourij, H.; Stechemesser, L.; Tilg, H.; Trauner, M.; Wagner, M.; Hofer, H.; Kiefer, F. W.; Fasching, P.; Roden, M. Austrian Multisociety Consensus on Metabolic Dysfunction-Associated Steatotic Liver Disease: Austrian Society of Gastroenterology and Hepatology (ÖGGH), Austrian Society of Diabetology (ÖDG), Austrian Society of Obesity (ÖAG). Wien. Klin. Wochenschr. 2025, 137 (S10), 307–319. 10.1007/s00508-025-02617-4.

(53) Uto, H.; Kanmura, S.; Takami, Y.; Tsubouchi, H. Clinical Proteomics for Liver Disease: A Promising Approach for Discovery of Novel Biomarkers. Proteome Sci. 2010, 8 (1), 70. 10.1186/1477-5956-8-70.

(54) Listopad, S.; Magnan, C.; Day, L. Z.; Asghar, A.; Stolz, A.; Tayek, J. A.; Liu, Z.-X.; Jacobs, J. M.; Morgan, T. R.; Norden-Krichmar, T. M. Identification of Integrated Proteomics and Transcriptomics Signature of Alcohol-Associated Liver Disease Using Machine Learning. *PLOS Digit*. Health 2024, 3 (2), e0000447. 10.1371/journal.pdig.0000447.

(55) Willebrords, J.; Pereira, I. V. A.; Maes, M.; Crespo Yanguas, S.; Colle, I.; Van Den Bossche, B.; Da Silva, T. C.; De Oliveira, C. P. M. S.; Andraus, W.; Alves, V. A.; Cogliati, B.; Vinken, M. Strategies, Models and Biomarkers in Experimental Non-Alcoholic Fatty Liver Disease Research. Prog. Lipid Res. 2015, 59, 106–125. 10.1016/j.plipres.2015.05.002.

(56) Klöting, N.; Fasshauer, M.; Dietrich, A.; Kovacs, P.; Schön, M. R.; Kern, M.; Stumvoll, M.; Blüher, M. Insulin-Sensitive Obesity. Am. J. Physiol.-Endocrinol. Metab. 2010, 299 (3), E506–E515. 10.1152/ajpendo.00586.2009.

(57) Anwar, M. Y.; Highland, H.; Buchanan, V. L.; Graff, M.; Young, K.; Taylor, K. D.; Tracy, R. P.; Durda, P.; Liu, Y.; Johnson, C. W.; Aguet, F.; Ardlie, K. G.; Gerszten, R. E.; Clish, C. B.; Lange, L. A.; Ding, J.; Goodarzi, M. O.; Chen, Y. I.; Peloso, G. M.; Guo, X.; Stanislawski, M. A.; Rotter, J. I.; Rich, S. S.; Justice, A. E.; Liu, C.; North, K. Machine Learning-based Clustering Identifies Obesity Subgroups with Differential Multi-omics Profiles and Metabolic Patterns. Obesity 2024, 32 (11), 2024–2034. 10.1002/oby.24137.

(58) Tanabe, H.; Sato, M.; Miyake, A.; Shimajiri, Y.; Ojima, T.; Narita, A.; Saito, H.; Tanaka, K.; Masuzaki, H.; Kazama, J. J.; Katagiri, H.; Tamiya, G.; Kawakami, E.; Shimabukuro, M. Machine Learning-Based Reproducible Prediction of Type 2 Diabetes Subtypes. Diabetologia 2024, 67 (11), 2446–2458. 10.1007/s00125-024-06248-8.

(59) Kopitar, L.; Kocbek, P.; Cilar, L.; Sheikh, A.; Stiglic, G. Early Detection of Type 2 Diabetes Mellitus Using Machine Learning-Based Prediction Models. Sci. Rep. 2020, 10 (1), 11981. 10.1038/s41598-020-68771-z.

(60) Carrasco-Zanini, J.; Pietzner, M.; Lindbohm, J. V.; Wheeler, E.; Oerton, E.; Kerrison, N.; Simpson, M.; Westacott, M.; Drolet, D.; Kivimaki, M.; Ostroff, R.; Williams, S. A.; Wareham, N. J.; Langenberg, C. Proteomic Signatures for Identification of Impaired Glucose Tolerance. Nat. Med. 2022, 28 (11), 2293–2300. 10.1038/s41591-022-02055-z.

(61) Figarska, S. M.; Rigdon, J.; Ganna, A.; Elmståhl, S.; Lind, L.; Gardner, C. D.; Ingelsson, E. Proteomic Profiles before and during Weight Loss: Results from Randomized Trial of Dietary Intervention. Sci. Rep. 2020, 10 (1), 7913. 10.1038/s41598-020-64636-7.

(62) Garin-Shkolnik, T.; Rudich, A.; Hotamisligil, G. S.; Rubinstein, M. FABP4 Attenuates PPARγ and Adipogenesis and Is Inversely Correlated With PPARγ in Adipose Tissues. Diabetes 2014, 63 (3), 900–911. 10.2337/db13-0436.

(63) Piening, B. D.; Zhou, W.; Contrepois, K.; Röst, H.; Gu Urban, G. J.; Mishra, T.; Hanson, B. M.; Bautista, E. J.; Leopold, S.; Yeh, C. Y.; Spakowicz, D.; Banerjee, I.; Chen, C.; Kukurba, K.; Perelman, D.; Craig, C.; Colbert, E.; Salins, D.; Rego, S.; Lee, S.; Zhang, C.; Wheeler, J.; Sailani, M. R.; Liang, L.; Abbott, C.; Gerstein, M.; Mardinoglu, A.; Smith, U.; Rubin, D. L.; Pitteri, S.; Sodergren, E.; McLaughlin, T. L.; Weinstock, G. M.; Snyder, M. P. Integrative Personal Omics Profiles during Periods of Weight Gain and Loss. Cell Syst. 2018, 6 (2), 157–170.e8. 10.1016/j.cels.2017.12.013.

(64) Trojnar, M.; Patro-Małysza, J.; Kimber-Trojnar, Ż.; Leszczyńska-Gorzelak, B.; Mosiewicz, J. Associations between Fatty Acid-Binding Protein 4–A Proinflammatory Adipokine and Insulin Resistance, Gestational and Type 2 Diabetes Mellitus. Cells 2019, 8 (3), 227. 10.3390/cells8030227.

(65) Bouhajja, H.; Bougacha-Elleuch, N.; Lucas, N.; Legrand, R.; Marrakchi, R.; Kaveri, S. V.; Jamoussi, K.; Ayadi, H.; Abid, M.; Mnif-Feki, M.; Fetissov, S. O. Affinity Kinetics of Leptin-Reactive Immunoglobulins Are Associated with Plasma Leptin and Markers of Obesity and Diabetes. Nutr. Diabetes 2018, 8 (1), 32. 10.1038/s41387-018-0044-y.

(66) Thorand, B.; Zierer, A.; Baumert, J.; Meisinger, C.; Herder, C.; Koenig, W. Associations between Leptin and the Leptin / Adiponectin Ratio and Incident Type 2 Diabetes in Middle-aged Men and Women: Results from the MONICA / KORA Augsburg Study 1984–2002. Diabet. Med. 2010, 27 (9), 1004–1011. 10.1111/j.1464-5491.2010.03043.x.

(67) Beijer, K.; Nowak, C.; Sundström, J.; Ärnlöv, J.; Fall, T.; Lind, L. In Search of Causal Pathways in Diabetes: A Study Using Proteomics and Genotyping Data from a Cross-Sectional Study. Diabetologia 2019, 62 (11), 1998–2006. 10.1007/s00125-019-4960-8.

(68) Hedbacker, K.; Birsoy, K.; Wysocki, R. W.; Asilmaz, E.; Ahima, R. S.; Farooqi, I. S.; Friedman, J. M. Antidiabetic Effects of IGFBP2, a Leptin-Regulated Gene. Cell Metab. 2010, 11 (1), 11–22. 10.1016/j.cmet.2009.11.007.

(69) Zaghlool, S. B.; Halama, A.; Stephan, N.; Gudmundsdottir, V.; Gudnason, V.; Jennings, L. L.; Thangam, M.; Ahlqvist, E.; Malik, R. A.; Albagha, O. M. E.; Abou-Samra, A. B.; Suhre, K. Metabolic and Proteomic Signatures of Type 2 Diabetes Subtypes in an Arab Population. Nat. Commun. 2022, 13 (1), 7121. 10.1038/s41467-022-34754-z.

(70) Crasan, I.-M.; Tanase, M.; Delia, C. E.; Gradisteanu-Pircalabioru, G.; Cimpean, A.; Ionica, E. Metaflammation’s Role in Systemic Dysfunction in Obesity: A Comprehensive Review. Int. J. Mol. Sci. 2025, 26 (21), 10445. 10.3390/ijms262110445.

(71) Chen, Z.-Z.; Gao, Y.; Keyes, M. J.; Deng, S.; Mi, M.; Farrell, L. A.; Shen, D.; Tahir, U. A.; Cruz, D. E.; Ngo, D.; Benson, M. D.; Robbins, J. M.; Correa, A.; Wilson, J. G.; Gerszten, R. E. Protein markers of diabetes discovered in an African American cohort. American Diabetes Association. 10.2337/figshare.21830886.

(72) Pan, Y.; Si, J. Endothelial Metabolic Reprogramming Links Diabetes to Atherosclerosis. Diabetes Metab. Syndr. Obes. 2026, *Volume* 19, 1–22. 10.2147/DMSO.S565805.

(73) Fejza, A.; Carobolante, G.; Poletto, E.; Camicia, L.; Schinello, G.; Di Siena, E.; Ricci, G.; Mongiat, M.; Andreuzzi, E. The Entanglement of Extracellular Matrix Molecules and Immune Checkpoint Inhibitors in Cancer: A Systematic Review of the Literature. Front. Immunol. 2023, 14, 1270981. 10.3389/fimmu.2023.1270981.

(74) Newman, A. C.; Nakatsu, M. N.; Chou, W.; Gershon, P. D.; Hughes, C. C. W. The Requirement for Fibroblasts in Angiogenesis: Fibroblast-Derived Matrix Proteins Are Essential for Endothelial Cell Lumen Formation. Mol. Biol. Cell 2011, 22 (20), 3791–3800. 10.1091/mbc.e11-05-0393.

(75) Ban, D.; Hua, S.; Zhang, W.; Shen, C.; Miao, X.; Liu, W. Costunolide Reduces Glycolysis-Associated Activation of Hepatic Stellate Cells via Inhibition of Hexokinase-2. Cell. Mol. Biol. Lett. 2019, 24 (1), 52. 10.1186/s11658-019-0179-4.

(76) Lian, N.; Jin, H.; Zhang, F.; Wu, L.; Shao, J.; Lu, Y.; Zheng, S. Curcumin Inhibits Aerobic Glycolysis in Hepatic Stellate Cells Associated with Activation of Adenosine Monophosphate-activated Protein Kinase. IUBMB Life 2016, 68 (7), 589–596. 10.1002/iub.1518.

(77) Jarvis, H.; Craig, D.; Barker, R.; Spiers, G.; Stow, D.; Anstee, Q. M.; Hanratty, B. Metabolic Risk Factors and Incident Advanced Liver Disease in Non-Alcoholic Fatty Liver Disease (NAFLD): A Systematic Review and Meta-Analysis of Population-Based Observational Studies. PLOS Med. 2020, 17 (4), e1003100. 10.1371/journal.pmed.1003100.

(78) Giraudi, P. J.; Salvoza, N.; Bonazza, D.; Saitta, C.; Lombardo, D.; Casagranda, B.; De Manzini, N.; Pollicino, T.; Raimondo, G.; Tiribelli, C.; Palmisano, S.; Rosso, N. Ficolin-2 Plasma Level Assesses Liver Fibrosis in Non-Alcoholic Fatty Liver Disease. Int. J. Mol. Sci. 2022, 23 (5), 2813. 10.3390/ijms23052813.

(79) Quek, J.; Chan, K. E.; Wong, Z. Y.; Tan, C.; Tan, B.; Lim, W. H.; Tan, D. J. H.; Tang, A. S. P.; Tay, P.; Xiao, J.; Yong, J. N.; Zeng, R. W.; Chew, N. W. S.; Nah, B.; Kulkarni, A.; Siddiqui, M. S.; Dan, Y. Y.; Wong, V. W.-S.; Sanyal, A. J.; Noureddin, M.; Muthiah, M.; Ng, C. H. Global Prevalence of Non-Alcoholic Fatty Liver Disease and Non-Alcoholic Steatohepatitis in the Overweight and Obese Population: A Systematic Review and Meta-Analysis. Lancet Gastroenterol. Hepatol. 2023, 8 (1), 20–30. 10.1016/S2468-1253(22)00317-X.

(80) Askeland, A.; Rasmussen, R. W.; Gjela, M.; Frøkjær, J. B.; Højlund, K.; Mellergaard, M.; Handberg, A. Non-Invasive Liver Fibrosis Markers Are Increased in Obese Individuals with Non-Alcoholic Fatty Liver Disease and the Metabolic Syndrome. Sci. Rep. 2025, 15 (1), 10652. 10.1038/s41598-025-85508-y.

(81) Finnson, K. W.; Philip, A. Endoglin in Liver Fibrosis. J. Cell Commun. Signal. 2012, 6 (1), 1–4. 10.1007/s12079-011-0154-y.

(82) Gavcar, D. C.; Danış, N.; Akca, M.; Döngelli, H.; Arayıcı, M.; Kırca, N. D.; Kızıldağ, S.; Akarsu, M. Noninvasive Fibrosis Triage in MASLD: Diagnostic Performance of Endocan and Endoglin (Cross-Sectional Study). Postgrad. Med. 2026, 138 (1), 44–52. 10.1080/00325481.2026.2630433.

(83) Vu, H.; Sun, Y.; Xiong, Z.; Tan, X.; Radford-Smith, D.; Causer, A.; Dickens, A. M.; Hyötyläinen, T.; Evstafev, I.; Oresic, M.; Nefzger, C.; O’Sullivan, E. D.; Watt, M. J.; Ramm, G. A.; Clouston, A.; Irvine, K. M.; Nguyen, Q. H.; Powell, E. E. Progressive Fibrosis in Human MASLD Is Associated with Spatially Linked Transcriptomic Signatures of Metabolic Reprogramming and Senescence. JHEP Rep. 2026, 8 (2), 101657. 10.1016/j.jhepr.2025.101657.

(84) Siddiqui, K.; George, T. P.; Mujammami, M.; Isnani, A.; Alfadda, A. A. The Association of Cell Adhesion Molecules and Selectins (VCAM-1, ICAM-1, E-Selectin, L-Selectin, and P-Selectin) with Microvascular Complications in Patients with Type 2 Diabetes: A Follow-up Study. Front. Endocrinol. 2023, 14, 1072288. 10.3389/fendo.2023.1072288.

(85) Yilmaz, O.; Erinc, O.; Gungordu, A. G.; Erdogan, M.; Algemi, M.; Akarsu, M. The Relationship of the Plasma Glycated CD59 Level with Microvascular Complications in Diabetic Patients and Its Evaluation as a Predictive Marker. J. Clin. Med. 2025, 14 (13), 4588. 10.3390/jcm14134588.

(86) Aydin Ozgur, B.; Coskunpinar, E.; Bilgic Gazioglu, S.; Yilmaz, A.; Musteri Oltulu, Y.; Cakmakoglu, B.; Deniz, G.; Gurol, A.; Yilmaz, M. Effects of Complement Regulators and Chemokine Receptors in Type 2 Diabetes. Immunol. Invest. 2021, 50 (5), 478–491. 10.1080/08820139.2020.1778022.

(87) Zhu, X.; Liang, F.; Yin, J.; Li, X.; Jiang, L.; Gao, Y.; Lu, Y.; Hu, Y.; Dai, N.; Su, J.; Yang, Z.; Yao, M.; Xiao, Y.; Ge, W.; Zhang, Y.; Zhong, Y.; Zhang, J.; Wu, M. Duration-Specific Association between Plasma IGFBP7 Levels and Diabetic Complications in Patients with Type 2 Diabetes Mellitus. Growth Horm. IGF Res. 2024, 75, 101574. 10.1016/j.ghir.2024.101574.

(88) Bracun, V.; Van Essen, B.; Voors, A. A.; Van Veldhuisen, D. J.; Dickstein, K.; Zannad, F.; Metra, M.; Anker, S.; Samani, N. J.; Ponikowski, P.; Filippatos, G.; Cleland, J. G. F.; Lang, C. C.; Ng, L. L.; Shi, C.; De Wit, S.; Aboumsallem, J. P.; Meijers, W. C.; Klip, Ij. T.; Van Der Meer, P.; De Boer, R. A. Insulin-Like Growth Factor Binding Protein 7 (IGFBP7), a Link Between Heart Failure and Senescence. ESC Heart Fail. 2022, 9 (6), 4167–4176. 10.1002/ehf2.14120.

(89) Lu, S.; Purohit, S.; Sharma, A.; Zhi, W.; He, M.; Wang, Y.; Li, C.-J.; She, J.-X. Serum Insulin-like Growth Factor Binding Protein 6 (IGFBP6) Is Increased in Patients with Type 1 Diabetes and Its Complications.

(90) Wu, W.; Xu, H.; Meng, Z.; Zhu, J.; Xiong, S.; Xia, X.; Lei, H. Axl Is Essential for In-Vitro Angiogenesis Induced by Vitreous From Patients With Proliferative Diabetic Retinopathy. Front. Med. 2021, 8, 787150. 10.3389/fmed.2021.787150.

