## Supplementary Figure 1 for "Machine learning uncovers circulating biomarkers and molecular heterogeneity in obesity and type 2 diabetes"

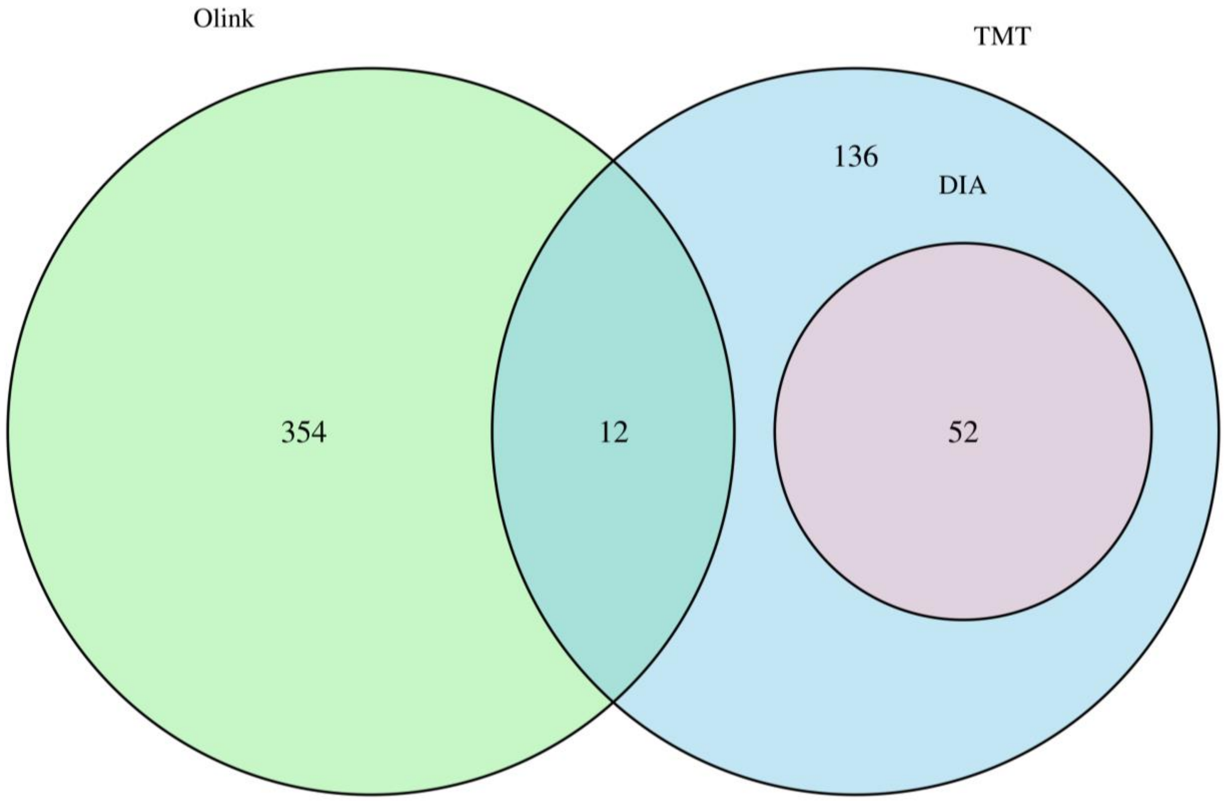


**Supplementary Figure 1. Integration of proteomics datasets obtained by Olink, TMT, and DIA platforms.** Venn diagram illustrating the overlap of quantified proteins across the Olink and TMT datasets, with DIA proteins represented within the TMT dataset. Due to data completeness across platforms, the Olink dataset was used for downstream analyses.
