## Supplementary Figure 2 for "Machine learning uncovers circulating biomarkers and molecular heterogeneity in obesity and type 2 diabetes"

**A.**

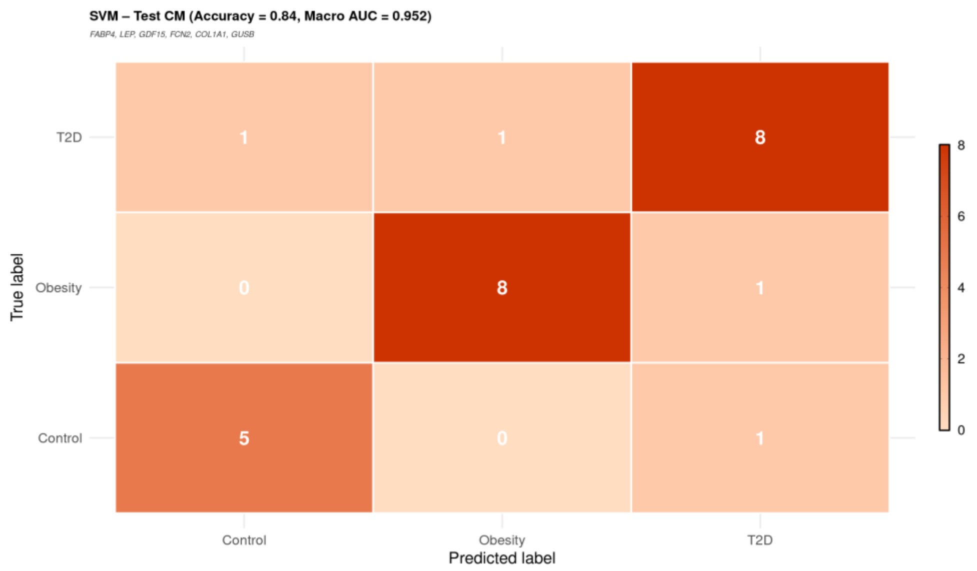


**B.**

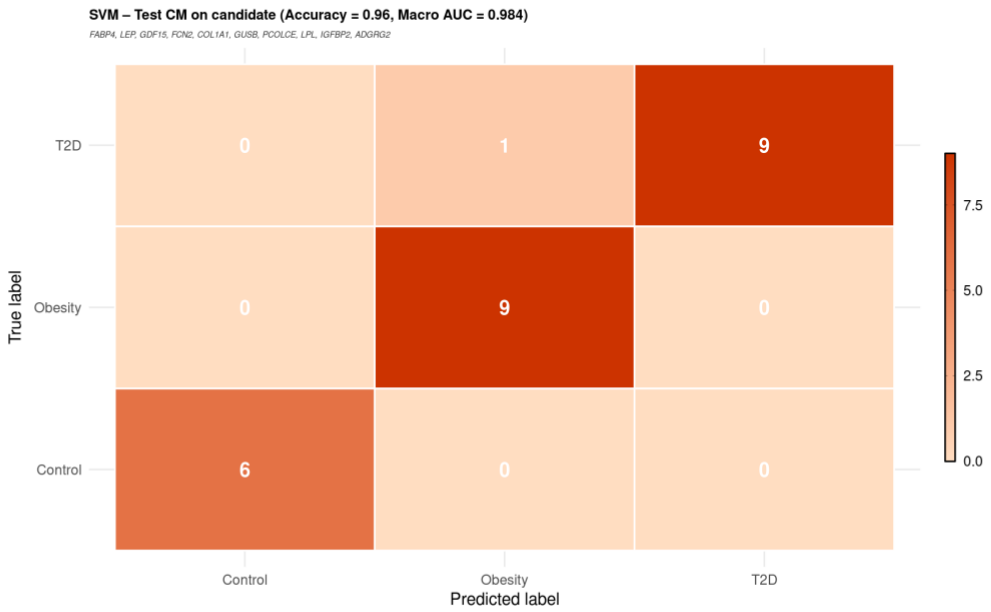


**Supplementary Figure 2. Performance of SVM classifiers using selected proteomic features for Control, Obesity, and T2D groups.** (A) Confusion matrix and performance metrics of the SVM model trained on six selected features (FABP4, LEP, GDF15, FCN2, COL1A1, GUSB). (B) Confusion matrix and performance metrics of the SVM model trained on ten selected features (FABP4, LEP, GDF15, FCN2, COL1A1, GUSB, PCOLCE, LPL, IGFBP2, ADGRG2). Rows represent true class labels and columns represent predicted labels.
