## Supplementary Figure 3 for "Machine learning uncovers circulating biomarkers and molecular heterogeneity in obesity and type 2 diabetes"

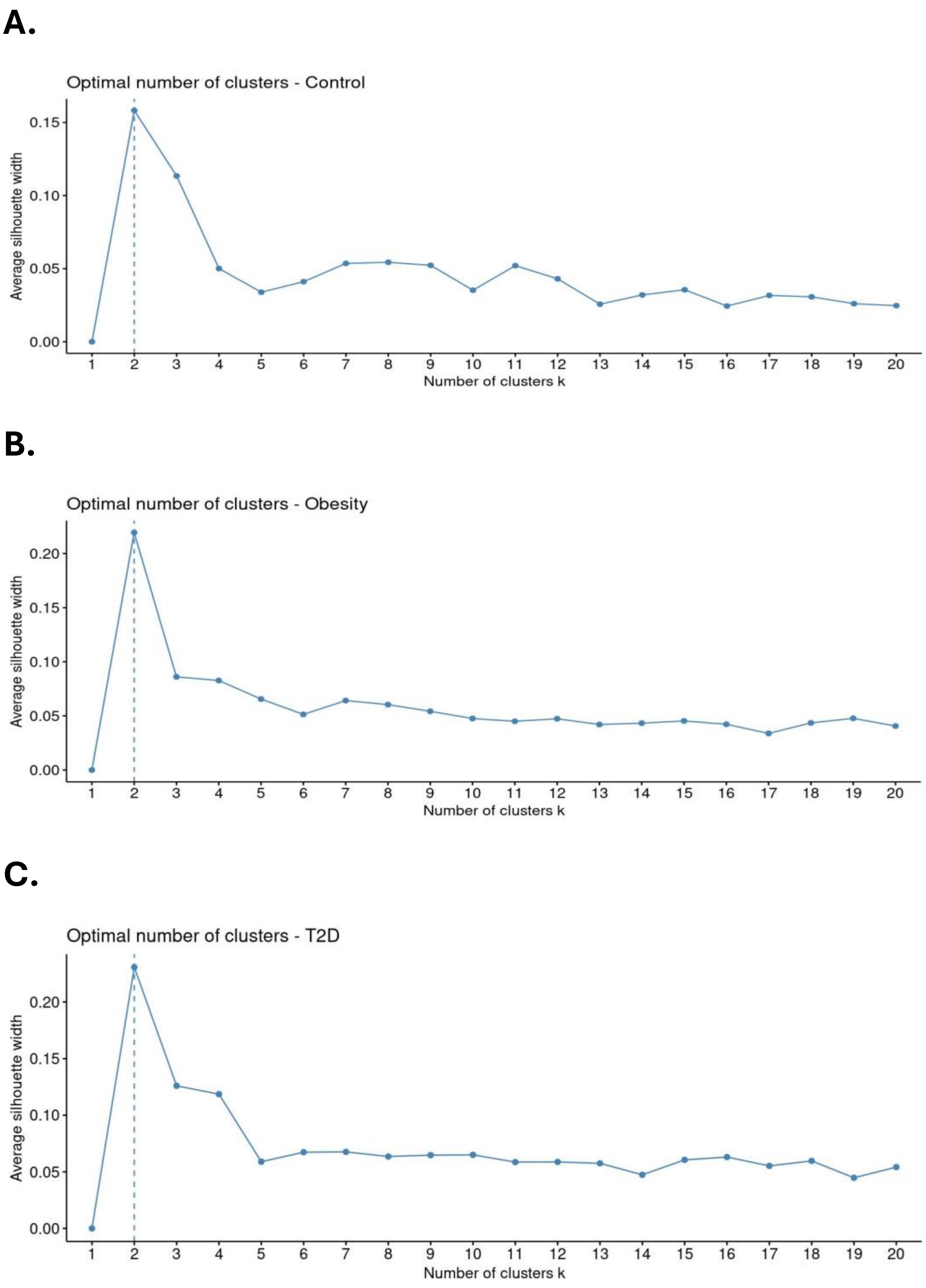


**Supplementary Figure 3. Determination of the optimal number of clusters using the silhouette method in Control, Obesity, and T2D groups.** (A-C) Average silhouette width was calculated across a range of cluster numbers (k = 1–20) for each group: (A) Control, (B) Obesity, and (C) Type 2 diabetes (T2D). In all three groups, the highest silhouette width was observed at k = 2, indicated by dashed vertical lines, suggesting that two clusters provide the most optimal partitioning of the samples. These results support the use of k = 2 for downstream clustering analyses in each group.
