## Supplementary Figure 4 for "Machine learning uncovers circulating biomarkers and molecular heterogeneity in obesity and type 2 diabetes"

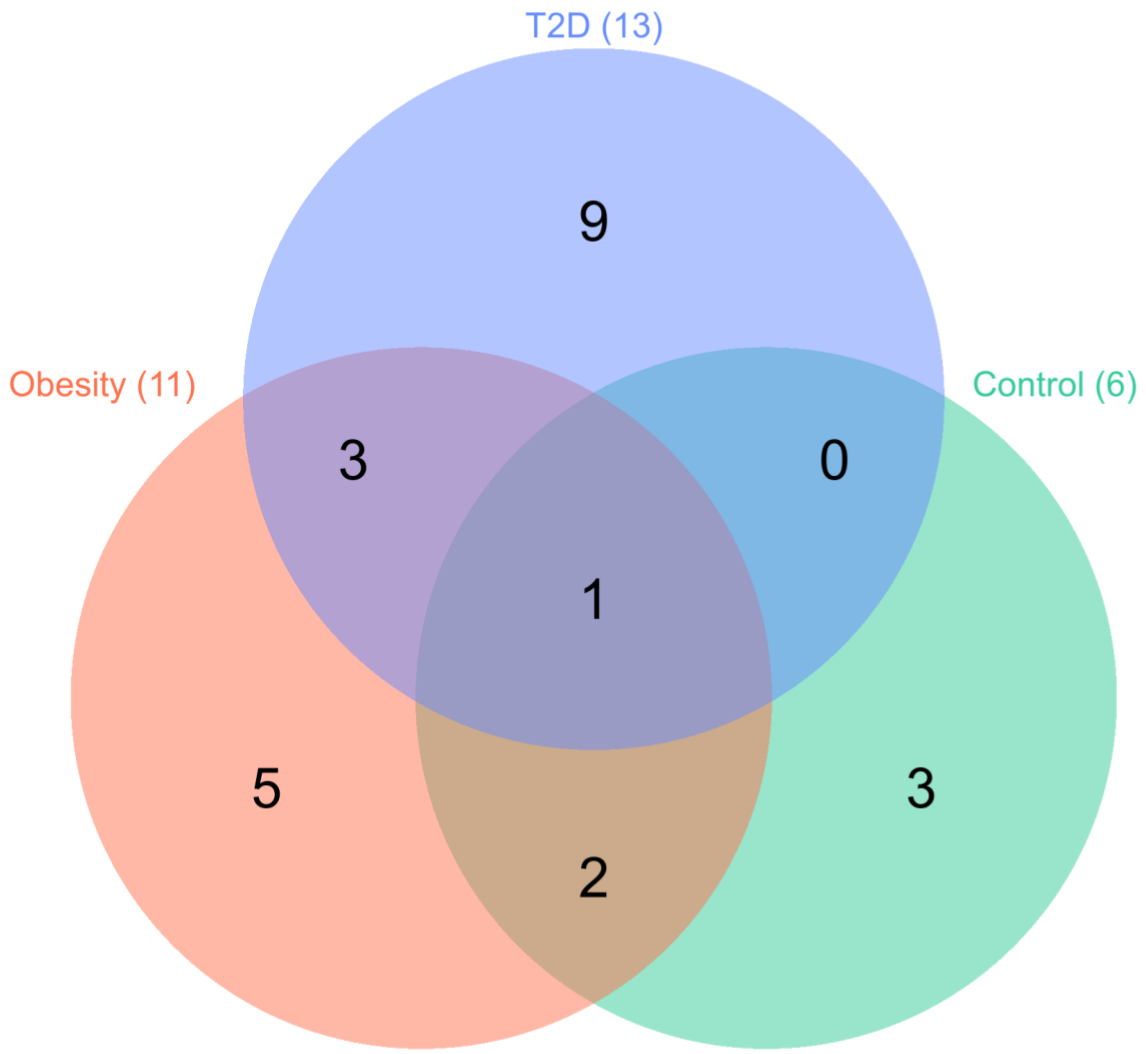


**Supplementary Figure 4. Cluster discriminating highly stable important features in Control, Obesity, and T2D groups.** A Venn diagram showing the distribution and overlap of proteins identified as highly stable important features distinguishing clusters within each group. Numbers in each circle indicate the count of proteins unique to or shared between groups.
