## Supplementary Figure 5 for "Machine learning uncovers circulating biomarkers and molecular heterogeneity in obesity and type 2 diabetes"

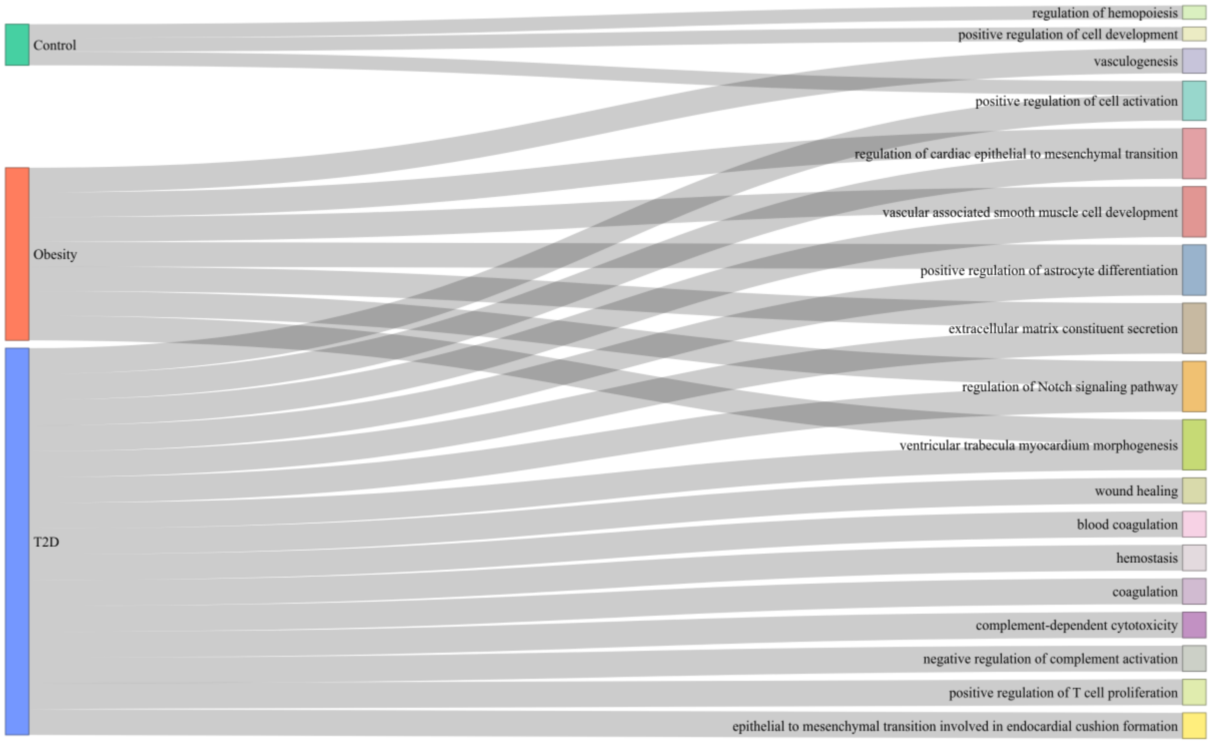


**Supplementary Figure 5. GO biological process enrichment of intragroup highly stable important features.** Sankey diagram showing the relationships between group-specific highly stable important features in Control, Obesity, and T2D groups and enriched Gene Ontology (GO) biological processes. Connections represent the association between proteins and significantly enriched GO terms.
