## Supplementary Figure 6 for "Machine learning uncovers circulating biomarkers and molecular heterogeneity in obesity and type 2 diabetes"

A.

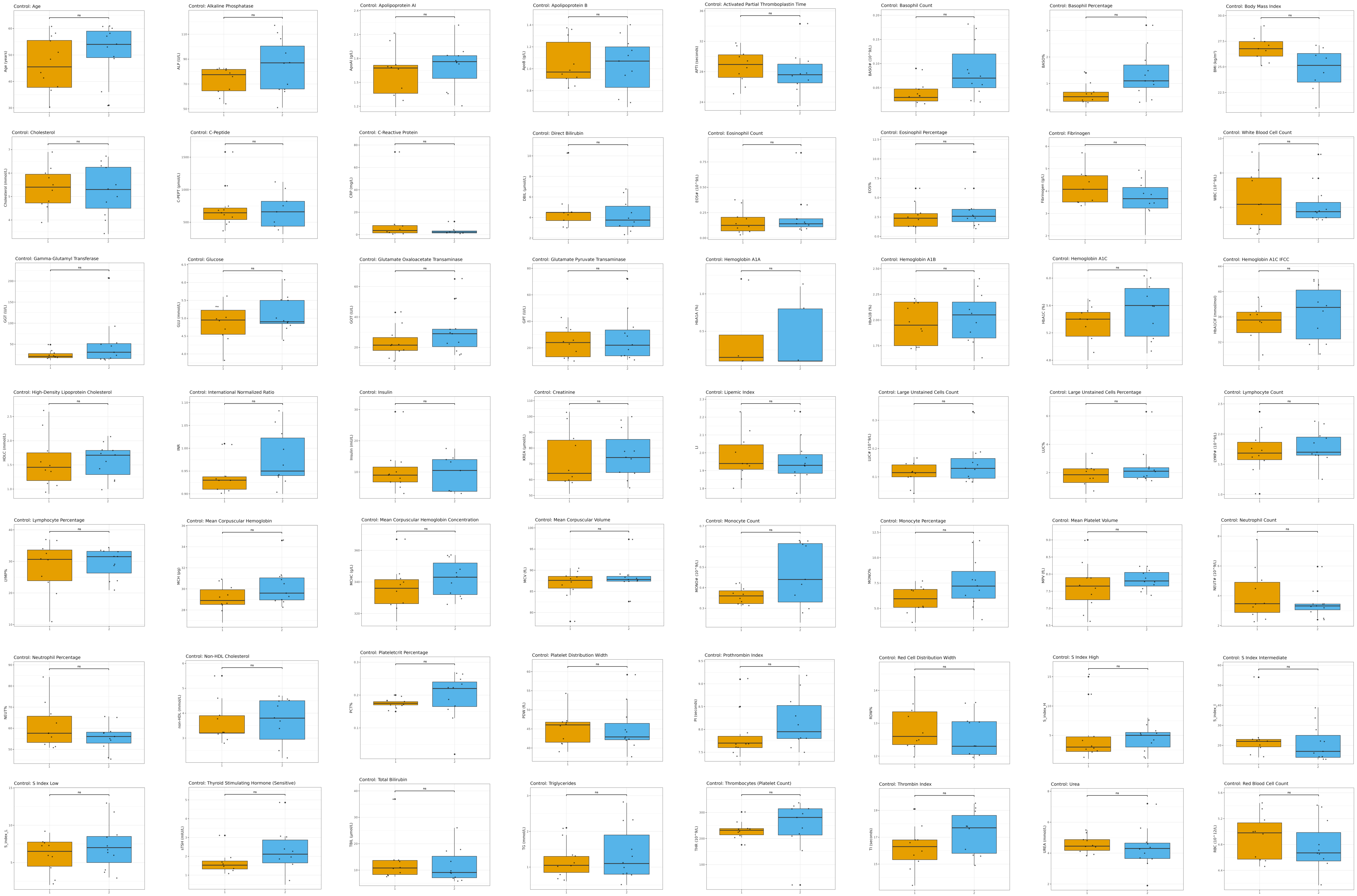
Control: Age
Control: Alkaline Phosphatase
Control: Apolipoprotein AI
Control: Apolipoprotein B
Control: Activated Partial Thromboplastin Time
Control: Basophil Count
Control: Basophil Percentage
Control: Body Mass Index
Control: Cholesterol
Control: C-Peptide
Control: C-Reactive Protein
Control: Direct Bilirubin
Control: Eosinophil Count
Control: Eosinophil Percentage
Control: Fibrinogen
Control: White Blood Cell Count
Control: Gamma-Glutamyl Transferase
Control: Glucose
Control: Glutamate Oxaloacetate Transaminase
Control: Glutamate Pyruvate Transaminase
Control: Hemoglobin A1A
Control: Hemoglobin A1B
Control: Hemoglobin A1C
Control: Hemoglobin A1C IFCC
Control: High-Density Lipoprotein Cholesterol
Control: International Normalized Ratio
Control: Insulin
Control: Creatinine
Control: Lipemic Index
Control: Large Unstained Cells Count
Control: Large Unstained Cells Percentage
Control: Lymphocyte Count
Control: Lymphocyte Percentage
Control: Mean Corpuscular Hemoglobin
Control: Mean Corpuscular Hemoglobin Concentration
Control: Mean Corpuscular Volume
Control: Monocyte Count
Control: Monocyte Percentage
Control: Mean Platelet Volume
Control: Neutrophil Count
Control: Neutrophil Percentage
Control: Non-HDL Cholesterol
Control: Plateletcrit Percentage
Control: Platelet Distribution Width
Control: Prothrombin Index
Control: Red Cell Distribution Width
Control: S Index High
Control: S Index Intermediate
Control: S Index Low
Control: Thyroid Stimulating Hormone (Sensitive)
Control: Total Bilirubin
Control: Triglycerides
Control: Thrombocytes (Platelet Count)
Control: Thrombin Index
Control: Urea
Control: Red Blood Cell Count

B.

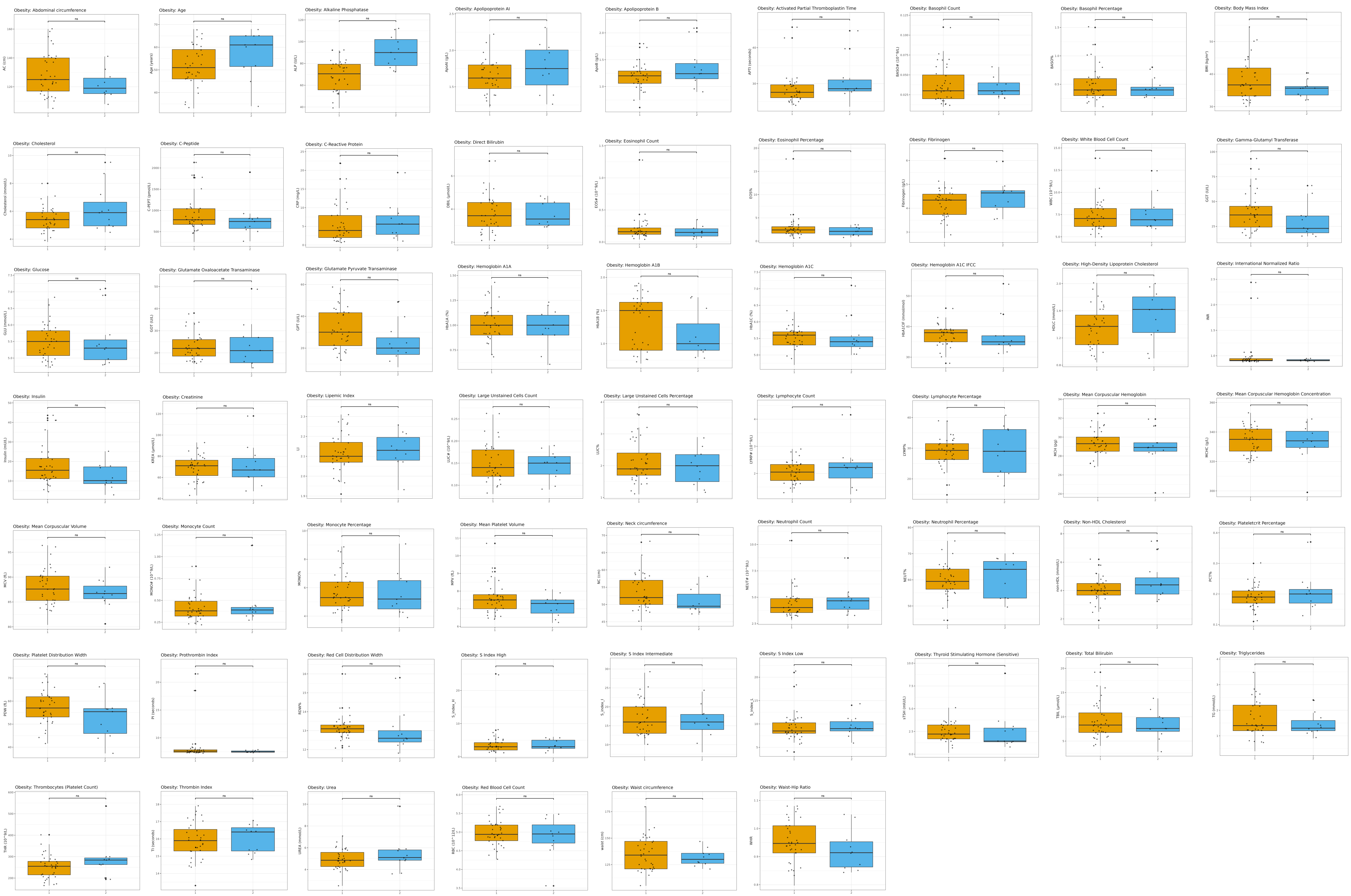
Obesity: Abdominal circumference
Obesity: Age
Obesity: Alkaline Phosphatase
Obesity: Apolipoprotein AI
Obesity: Apolipoprotein B
Obesity: Activated Partial Thromboplastin Time
Obesity: Basophil Count
Obesity: Basophil Percentage
Obesity: Body Mass Index
Obesity: Cholesterol
Obesity: C-Peptide
Obesity: C-Reactive Protein
Obesity: Direct Bilirubin
Obesity: Eosinophil Count
Obesity: Eosinophil Percentage
Obesity: Fibrinogen
Obesity: White Blood Cell Count
Obesity: Gamma-Glutamyl Transferase
Obesity: Glucose
Obesity: Glutamate Oxaloacetate Transaminase
Obesity: Glutamate Pyruvate Transaminase
Obesity: Hemoglobin A1A
Obesity: Hemoglobin A1B
Obesity: Hemoglobin A1C
Obesity: Hemoglobin A1C IFCC
Obesity: High-Density Lipoprotein Cholesterol
Obesity: International Normalized Ratio
Obesity: Insulin
Obesity: Creatinine
Obesity: Lipemic Index
Obesity: Large Unstained Cells Count
Obesity: Large Unstained Cells Percentage
Obesity: Lymphocyte Count
Obesity: Lymphocyte Percentage
Obesity: Mean Corpuscular Hemoglobin
Obesity: Mean Corpuscular Hemoglobin Concentration
Obesity: Mean Corpuscular Volume
Obesity: Monocyte Count
Obesity: Monocyte Percentage
Obesity: Mean Platelet Volume
Obesity: Neck circumference
Obesity: Neutrophil Count
Obesity: Neutrophil Percentage
Obesity: Non-HDL Cholesterol
Obesity: Plateletcrit Percentage
Obesity: Platelet Distribution Width
Obesity: Prothrombin Index
Obesity: Red Cell Distribution Width
Obesity: S Index High
Obesity: S Index Intermediate
Obesity: S Index Low
Obesity: Thyroid Stimulating Hormone (Sensitive)
Obesity: Total Bilirubin
Obesity: Triglycerides
Obesity: Thrombocytes (Platelet Count)
Obesity: Thrombin Index
Obesity: Urea
Obesity: Red Blood Cell Count
Obesity: Waist circumference
Obesity: Waist-Hip Ratio

C.

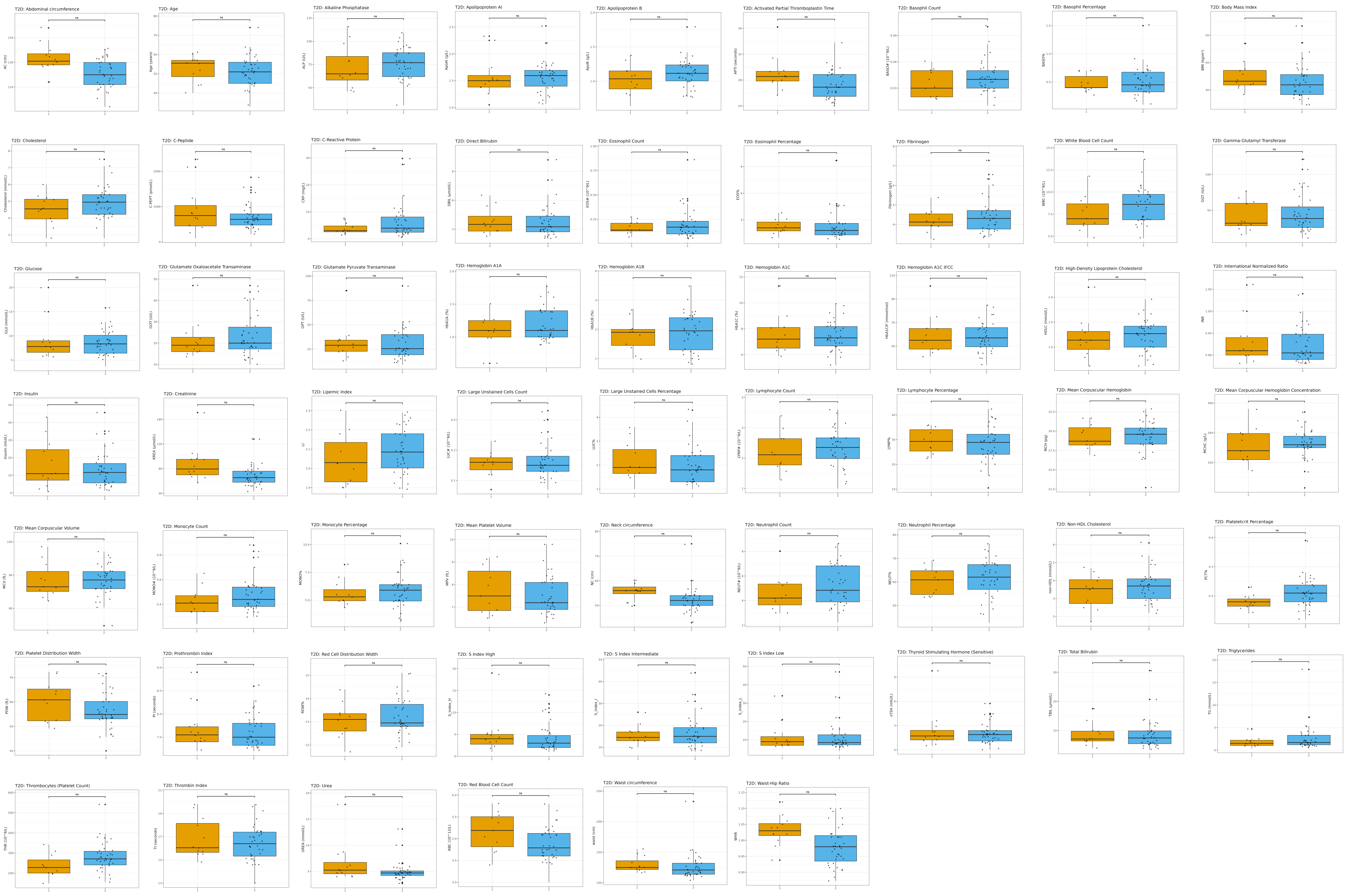
T2D: Abdominal circumference
T2D: Age
T2D: Alkaline Phosphatase
T2D: Apolipoprotein AI
T2D: Apolipoprotein B
T2D: Activated Partial Thromboplastin Time
T2D: Basophil Count
T2D: Basophil Percentage
T2D: Body Mass Index
T2D: Cholesterol
T2D: C-Peptide
T2D: C-Reactive Protein
T2D: Direct Bilirubin
T2D: Eosinophil Count
T2D: Eosinophil Percentage
T2D: Fibrinogen
T2D: White Blood Cell Count
T2D: Gamma-Glutamyl Transferase
T2D: Glucose
T2D: Glutamate Oxaloacetate Transaminase
T2D: Glutamate Pyruvate Transaminase
T2D: Hemoglobin A1A
T2D: Hemoglobin A1B
T2D: Hemoglobin A1C
T2D: Hemoglobin A1C IFCC
T2D: High-Density Lipoprotein Cholesterol
T2D: International Normalized Ratio
T2D: Insulin
T2D: Creatinine
T2D: Lipemic Index
T2D: Large Unstained Cells Count
T2D: Large Unstained Cells Percentage
T2D: Lymphocyte Count
T2D: Lymphocyte Percentage
T2D: Mean Corpuscular Hemoglobin
T2D: Mean Corpuscular Hemoglobin Concentration
T2D: Mean Corpuscular Volume
T2D: Monocyte Count
T2D: Monocyte Percentage
T2D: Mean Platelet Volume
T2D: Neck circumference
T2D: Neutrophil Count
T2D: Neutrophil Percentage
T2D: Non-HDL Cholesterol
T2D: Plateletcrit Percentage
T2D: Platelet Distribution Width
T2D: Prothrombin Index
T2D: Red Cell Distribution Width
T2D: S Index High
T2D: S Index Intermediate
T2D: S Index Low
T2D: Thyroid Stimulating Hormone (Sensitive)
T2D: Total Bilirubin
T2D: Triglycerides
T2D: Thrombocytes (Platelet Count)
T2D: Thrombin Index
T2D: Urea
T2D: Red Blood Cell Count
T2D: Waist circumference
T2D: Waist-Hip Ratio

**Supplementary Figure 6. Distribution of clinical variables between two clusters within each study group.** (A-C) Boxplots showing the distribution of clinical and laboratory variables between the two clusters identified in the Control (A, N = 56), Obesity (B, N = 60), and T2D (C, N = 60) groups. Each panel presents comparisons across multiple variables, with boxes representing the interquartile range (IQR), center lines indicating the median, and whiskers denoting the data range, and points representing individual observations. Statistical significance between clusters was assessed using the Wilcoxon rank-sum test.
